# Protective ApoE variants support neuronal function by extracting peroxidated lipids

**DOI:** 10.1101/2025.03.05.641722

**Authors:** Isha Ralhan, Alison D. Do, Ju-Young Bae, Femke M. Feringa, Wendy Cai, Jinlan Chang, Kennedi Chik, Nathanael YJ Lee, Chris Gerry, Rik van der Kant, Jesse Jackson, Emily L. Ricq, Maria S. Ioannou

## Abstract

ApoE mediates the transport of lipids from neurons to glial lipid droplets. ApoE4, a major risk factor for Alzheimer’s disease, impairs this transport pathway, increasing risk for neurodegeneration. ApoE2 and ApoE3 Christchurch (ApoE3Ch) variants confer resistance to developing the disease, yet little is known regarding how these protective variants affect lipid transport. Here, we explored how lipoprotein particles containing different ApoE isoforms affect neuronal health in vitro and in intact rodent hippocampi. We demonstrate that ApoE2 and ApoE3Ch particles protect neurons from ferroptosis by preferentially extracting peroxidated unsaturated lipids through the neuronal ABCA7 transporter. ApoE4 particles, on the other hand, exacerbate the effects of these toxic lipids leading to endolysosomal dysfunction. By reducing the peroxidated lipid burden in ApoE4 neurons, ApoE2 and ApoE3Ch particles rescue endolysosomal function and restore defects in neuronal activity. Our findings reveal a new mechanism by which ApoE2 and ApoE3Ch isoforms protect neurons from neurodegenerative disease.

**Highlights:**

- ApoE2 and ApoE3Ch reduce neuronal ferroptosis by peroxidated lipid efflux via ABCA7
- Lysosomal defects caused by lipid peroxidation are rescued by ApoE2 and ApoE3Ch
- ApoE2 and ApoE3Ch particles protect ApoE4 hippocampi from toxicity
- ApoE2 and ApoE3Ch particles rescue impaired neuronal activity in ApoE4 hippocampi

## INTRODUCTION

Alzheimer’s disease (AD) is a progressive neurodegenerative disease and the most common cause of dementia. Pathological hallmarks of the disease include amyloid-β (Aβ) plaques and dysregulated lipid homeostasis^1,2^. Lipid transport between neurons and glia is also essential for maintaining brain health. Failure to transport neuronal lipids to glia during oxidative stress results in neurodegeneration^3,4^. While the mechanisms of this lipid transport pathway are not fully understood, there is a clear role for Apolipoprotein E (ApoE).

ApoE is highly expressed in the brain and is the strongest genetic risk factor for late-onset AD^5,6^. There are three common isoforms, of which *APOE4* confers elevated risk for developing AD^5^, while *APOE2* is neuroprotective relative to the most common isoform *APOE3*^7^. ApoE functions as a component of high-density lipoprotein-like particles in the brain where it mediates intercellular lipid transport^8–10^. Consistently, ApoE2 and ApoE3 promote neuron-to-glia lipid transport and subsequent glial lipid droplet formation while ApoE4 is unable to do so^3^. This indicates a genotype-specific effect on lipid transport from neurons to glia^3^. ApoE also prevents neuronal ferroptosis^11^, a form of cell death that is implicated in AD and characterized by the accumulation of lipid peroxides^12^. This points to a failure in transporting peroxidated lipids from neurons to glia in disease, but how ApoE isoforms regulate the efflux of peroxidated lipids from neurons remains poorly understood.

According to the ApoE cascade hypothesis, the properties of ApoE isoforms differentially impact cellular function thereby triggering a cascade of events that promote AD pathology^13^. For example, the biochemical properties of ApoE4, which include altered lipid binding, precede defects in cellular function such as endolysosomal defects and Aβ pathology^14^. Interventions that target the early phase of the ApoE cascade are more likely to be effective against age-related cognitive decline in AD^13^, yet most of our knowledge of how ApoE contributes to AD is centered on the detrimental effects of ApoE4^13^. Much less is known regarding the role of the protective variants ApoE2 and ApoE3 Christchurch (ApoE3Ch)^15^ which are likely to reveal important insight into the disease mechanism. The protective effects of ApoE2 and ApoE3Ch have been recently linked to reduced receptor binding and lipid uptake in neurons and microglia^16^, but how these protective variants affect lipid efflux from neurons and the type of lipids being transported remains unclear.

Here, we investigate the mechanisms by which ApoE influences lipid export from neurons. We demonstrate that endocytosis of ApoE4 particles by neurons triggers the accumulation of peroxidated lipids, dysregulates the endolysosomal system and sensitizes these neurons to ferroptosis. However, this toxicity can be rescued by the addition of ApoE2 and ApoE3Ch particles which selectively efflux peroxidated unsaturated fatty acids through the ABCA7 transporter. In fact, these protective particles restore neuronal activity in ApoE4 hippocampi. They also rescue the lipid peroxidation and endolysosomal defects caused by ApoE4 genotype in rat neurons, human iPSC-derived neurons and ex vivo hippocampi. Together, our study reveals a mechanism by which ApoE2 and ApoE3Ch protect neurons from lipid-mediated toxicity associated with neurodegenerative disease.

## RESULTS

### ApoE2 and ApoE3Ch reduce lipid storage in neurons

While isoform specific effects of ApoE on lipid storage are better studied in glia, we sought to explore how ApoE isoforms affect neuronal lipids. Hippocampal rat neurons cultured from wild-type and human ApoE4 targeted replacement rats were treated with cytosine arabinoside to minimize contaminating glia (Figure S1A). We generated 1-palmitoyl-2-oleoyl-sn-glycero-3-phosphocholine particles with recombinant human ApoE2, ApoE3, ApoE4 or ApoE3Ch. The purity of ApoE protein in the particles was confirmed by SDS-PAGE and Coomassie staining (Figures S1B and S1C). Similar to those found in the brain^17,18^, ApoE particles were approximately 10-17 nm in diameter, as determined by native PAGE, transmission electron microscopy, and dynamic light scattering (Figures S1D-S1F; Table S1). We treated neurons with these particles and evaluated the number of BODIPY-493/503-positive lipid droplets as a measure of lipid storage. Both wild-type and ApoE4 neurons decreased lipid droplets with ApoE2, ApoE3 or ApoE3Ch particles, while ApoE4 particles had no effect (Figures S2A-S2F). These results suggest an isoform-dependent role for protective ApoE particles in removing lipids from neurons.

### ApoE2 and ApoE3Ch preferentially extract peroxidated unsaturated lipids from neurons

Since we observed an isoform-specific effect of ApoE particles on neuronal lipid accumulation, we hypothesized that ApoE2 and ApoE3Ch particles would be better at extracting fatty acids from neurons. To test this, neurons were first preloaded with BODIPY 558/568 (Red-C12), a fluorescently labelled saturated fatty acid, and the amount of Red-C12 released into the media was detected using a fluorescence plate reader. We found that both damaging and protective ApoE particles promoted Red-C12 efflux at similar levels (Figures 1A, and S3A-S3D). This indicates that the isoform-specific effects on neuronal lipid accumulation are mediated by mechanisms beyond saturated fatty acid efflux.

**Figure 1.**
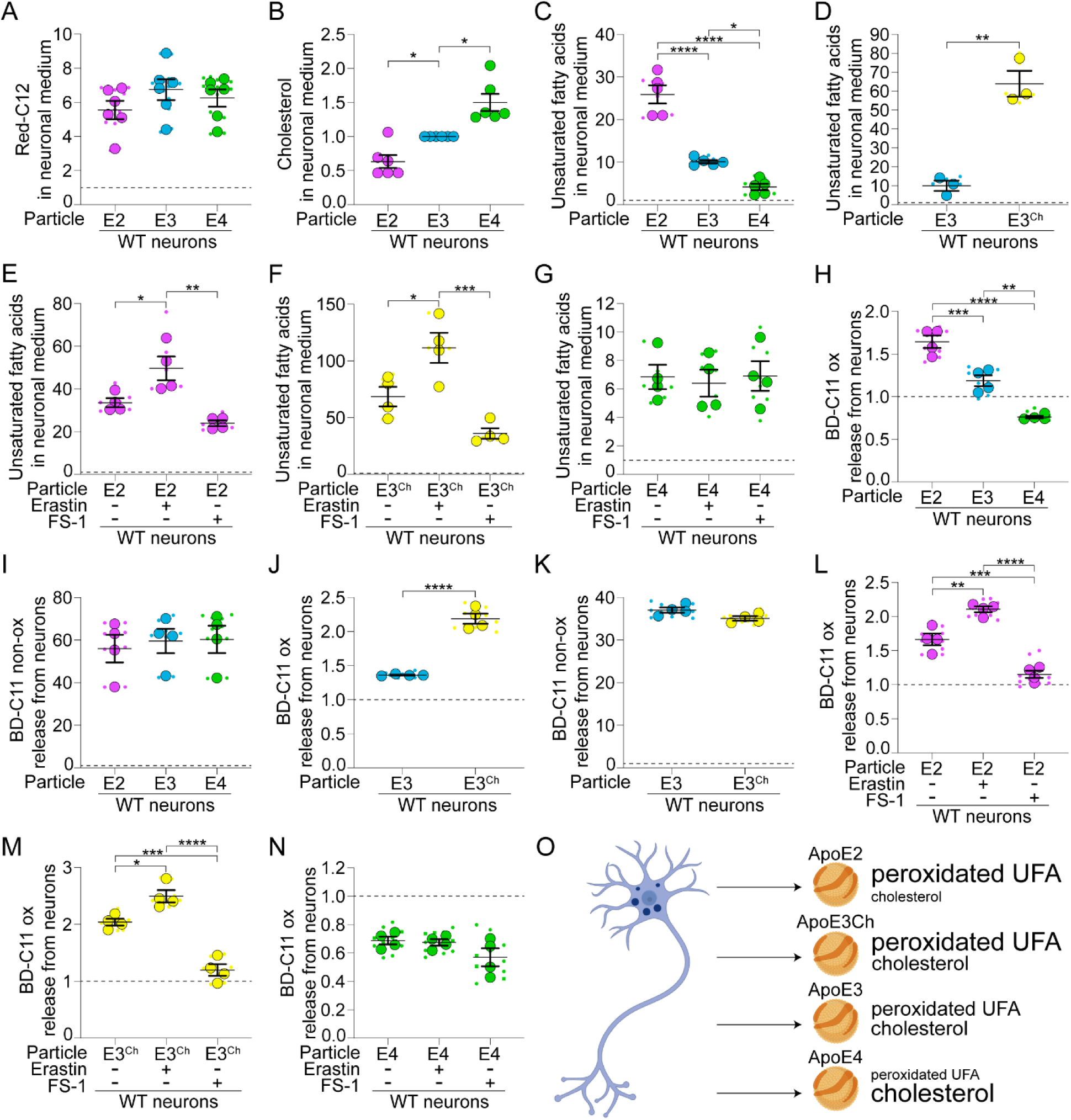
ApoE2 and ApoE3Ch particles extract peroxidated unsaturated fatty acids from neurons (A) WT neuron-conditioned HBSS following ApoE2, ApoE3 or ApoE4 particle treatment analyzed for Red-C12 fluorescence and normalized to non-particle treated control neurons (dashed line). n = 6 independent experiments; mean ± SEM. One-way ANOVA with Tukey’s post-test. (B) WT neuron-conditioned HBSS following ApoE2, ApoE3 or ApoE4 particle treatment analyzed for endogenous cholesterol and normalized to ApoE3 particle treated neurons. n = 6 independent experiments; mean ± SEM. One sample t test with Bonferroni correction. (C and D) WT neuron-conditioned HBSS following ApoE2, ApoE3, ApoE4 or ApoE3^Ch^ particle treatment analyzed for endogenous unsaturated fatty acids and normalized to non-particle treated control neurons (dashed line). (C) n = 5 independent experiments; mean ± SEM. One-way ANOVA with Tukey’s post-test. (D) n = 3 independent experiments; mean ± SEM. Two-tailed Student’s t test. (E–G) WT neuron-conditioned HBSS following erastin, ferrostatin-1 (FS-1), ApoE2, ApoE3, ApoE4 or ApoE3^Ch^ particle treatment analyzed for endogenous unsaturated fatty acids and normalized to DMSO treated control neurons without particles (dashed line). n = 4 independent experiments; mean ± SEM. One-way ANOVA with Tukey’s post-test. (H and I) WT neuron-conditioned HBSS following ApoE2, ApoE3 or ApoE4 particle treatment analyzed for peroxidated lipids (BD-C11 ox; H) and non-peroxidated lipids (BD-C11 non-ox; I). Particle treated neurons were normalized to non-particle treated control neurons (dashed line). N = 4 independent experiments; mean ± SEM. One-way ANOVA with Tukey’s post-test. (J and K) WT neuron-conditioned HBSS following ApoE3 or ApoE3^Ch^ particle treatment analyzed for peroxidated lipids (BD-C11 ox; J) and non-peroxidated lipids (BD-C11 non-ox; K). Particle treated neurons were normalized to non-particle treated control neurons (dashed line). N = 4 independent experiments; mean ± SEM. Two-tailed Student’s t test. (L-N) WT neuron-conditioned HBSS following ApoE2, ApoE3^Ch^ or ApoE4 particle treatment with or without erastin or FS-1, analyzed for peroxidated lipids (BD-C11 ox) and normalized to DMSO treated control neurons without particles (dashed line). n = 4 independent experiments; mean ± SEM. One-way ANOVA with Tukey’s post-test. (O) Schematic of ApoE-mediated lipid efflux from neurons. For all graphs independent replicates are in large shapes and technical replicates in small shapes; *p < 0.05, **p < 0.01, ***p < 0.001, ****p < 0.0001.

We wondered if instead ApoE isoforms differentially extracted other lipids from neurons, such as cholesterol or unsaturated fatty acids. Neurons were preloaded with cholesterol fluorescently tagged with nitrobenzoxadiazole (NBD) and the release of NBD-cholesterol in the media was detected using a fluorescence plate reader. All ApoE particles increased NBD-cholesterol in the medium with ApoE4 particles being more effective in the targeted replacement neurons (Figures S3E-S3H). To confirm these results, we measured endogenous cholesterol. Again, ApoE4 extracted the most cholesterol from neurons, ApoE2 extracted the least, while ApoE3Ch was equal to that of ApoE3 (Figures 1B and S3I). While this phenotype is consistent with increased cholesterol in ApoE4 astrocytes^19^, it still cannot account for how ApoE particles reduce neutral lipid accumulation. We next tested how ApoE particles affect the release of endogenous unsaturated fatty acids from neurons. Strikingly, protective ApoE2 and ApoE3Ch particles stimulated robust unsaturated fatty acid efflux, while ApoE4 particles were much less effective (Figures 1C and 1D). This indicates an isoform-dependent switch in the type of lipids being extracted from neurons.

Since unsaturated fatty acids are susceptible to lipid peroxidation^12^, we next asked whether the release of unsaturated fatty acids is affected by peroxidation. We analyzed unsaturated fatty acid efflux from neurons treated with erastin or ferrostatin-1, to increase or decrease lipid peroxidation, respectively. Both ApoE2 and ApoE3Ch particles extracted more unsaturated fatty acids when lipid peroxidation was increased and a slight but non-significant decrease when lipid peroxidation was reduced (Figures 1E and 1F). Lipid peroxidation did not affect the amount of unsaturated fatty acids extracted by ApoE4 (Figure 1G). As a second strategy, we incubated neurons with the lipid peroxidation sensor BODIPY 581/591 (BD-C11). This sensor incorporates into cellular membranes and shifts its fluorescence emission from 590 nm to 510 nm following peroxidation of a diene that behaves as an unsaturated fatty acyl moiety. Consistently, we found that ApoE2 and ApoE3Ch particles promoted peroxidated BD-C11 efflux, with no difference in non-peroxidated BD-C11 released (Figures 1H-1K). Conversely, ApoE4 particles failed to extract peroxidated BD-C11 (Figure 1H). Next, we repeated this experiment in the presence of erastin or ferrostatin-1 to modulate lipid peroxide levels. Again, ApoE2 and ApoE3Ch particles extracted more peroxidated BD-C11 from neurons treated with erastin and less peroxidated BD-C11 from neurons treated with ferrostatin-1 (Figures 1L, 1M, S3J and S3K). Modulating lipid hydroperoxide levels had no effect on BD-C11 efflux in the presence of ApoE4 particles (Figures 1N and S3L).

Collectively, these data reveal an isoform-specific switch in the ability of ApoE to extract different lipids from neurons. Neuroprotective ApoE2 and ApoE3Ch preferentially extract peroxidated unsaturated fatty acids, whereas ApoE4 lacks this capacity and prefers cholesterol (Figure 1O).

### ApoE2 and ApoE3Ch particles prevent while ApoE4 particles promote neuronal ferroptosis

Since lipid peroxidation increases unsaturated lipid efflux by protective ApoE particles, we reasoned that this would relieve neurons of lipid peroxidation. To test this, we stained neurons with BD-C11 and imaged the shift in fluorescence emission peak from 590 nm to 510 nm upon lipid peroxidation, using microscopy. As expected, erastin increased lipid peroxidation as evidenced by a higher BD-C11 ratio (510/590 nm) (Figures 2A and 2B). This increase in lipid peroxidation was rescued by the addition of ApoE2 and ApoE3 but not ApoE4 particles (Figures 2A and 2B).

**Figure 2.**
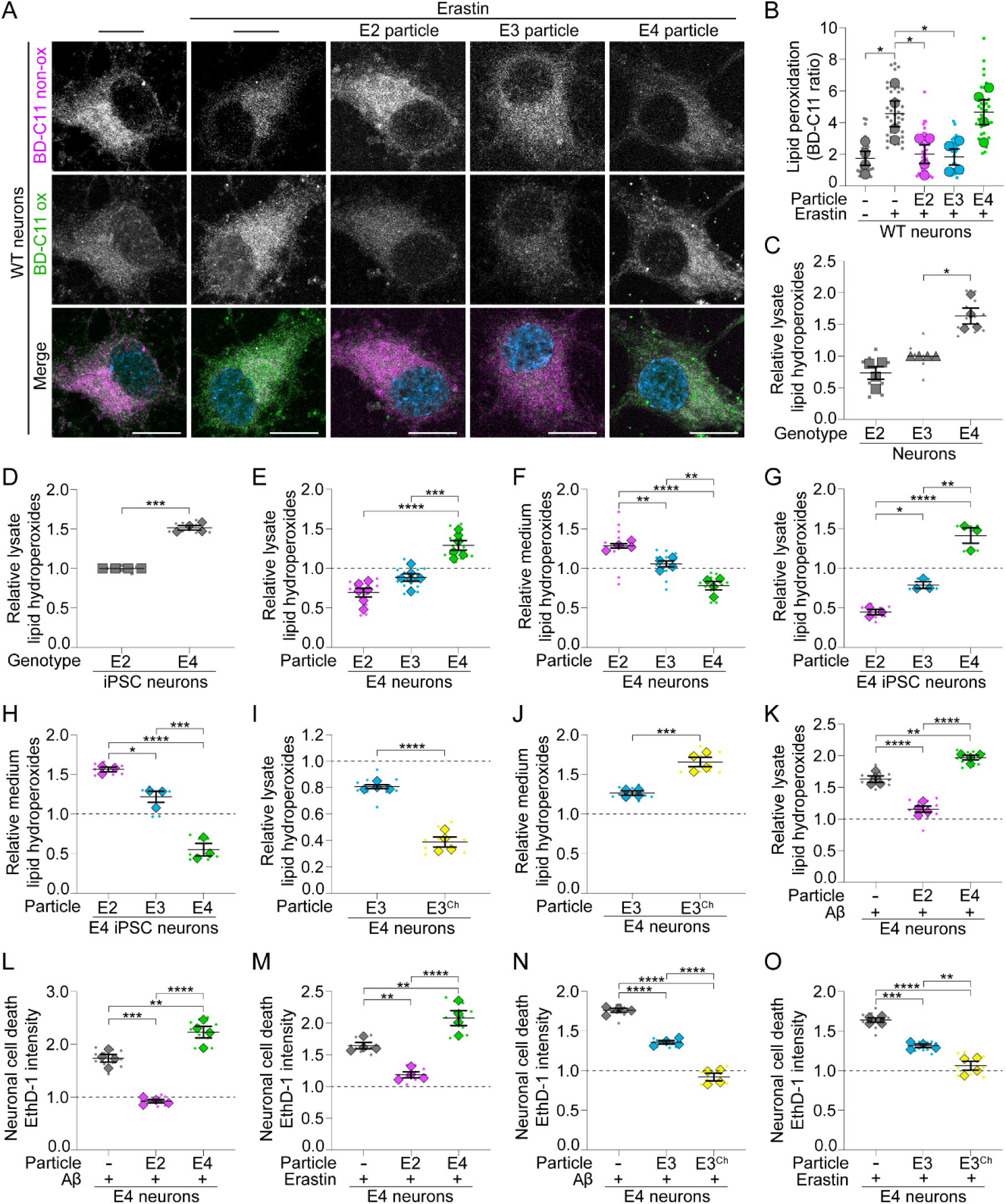
ApoE2 and ApoE3Ch protect neurons from lipid peroxidation mediated-toxicity (A and B) Airyscan images of WT neurons displayed as maximum intensity projections following treatment in media with ApoE2, ApoE3, or ApoE4 particles and/or erastin, labeled with BD-C11. Emission at 591 nm and 488 nm indicate non-peroxidated (BD-C11 non-ox) and peroxidated (BD-C11 ox) lipids, respectively. n = 4 independent experiments; mean ± SEM. One-way ANOVA with Dunnett’s post-test. Scale bars are 10 μm. (C) ApoE targeted replacement neuronal lysates analyzed for lipid hydroperoxides and normalized to ApoE3 neurons following HBSS treatment. n = 4 independent experiments; mean ± SEM. One sample t test with Bonferroni correction. (D) ApoE2 and ApoE4 human induced pluripotent stem cell (iPSC) neurons analyzed for lipid hydroperoxides and normalized to ApoE2 neurons following HBSS treatment. n = 4 independent experiments; mean ± SEM. One sample t test with Bonferroni correction. (E and F) ApoE4 neuronal lysates (E) and neuron-conditioned HBSS (F) following ApoE2, ApoE3 or ApoE4 particle treatment, analyzed for lipid hydroperoxides and normalized to non-particle treated control neurons (dashed line). n = 4-6 independent experiments; mean ± SEM. One-way ANOVA with Tukey’s post-test. (G and H) ApoE4 iPSC neuronal lysates (G) and neuron-conditioned HBSS (H) following ApoE2, ApoE3 or ApoE4 particle treatment, analyzed for lipid hydroperoxides and normalized to non-particle treated control neurons (dashed line). n = 3 independent experiments; mean ± SEM. One-way ANOVA with Tukey’s post-test. (I and J) ApoE4 neuronal lysates (I) and neuron-conditioned HBSS (J) following ApoE3 or ApoE3^Ch^ particle treatment, analyzed for lipid hydroperoxides and normalized to non-particle treated control neurons (dashed line). n = 4 independent experiments; mean ± SEM. Two-tailed Student’s t test. (K) ApoE4 neuronal lysates following treatment in media with Aβ and ApoE2 or ApoE4 particles, analyzed for lipid hydroperoxides and normalized to NaOH treated control neurons (dashed line). n = 4 independent experiments; mean ± SEM. One-way ANOVA with Tukey’s post-test. (L-O) ApoE4 neurons following treatment in media with Aβ or erastin and ApoE2, ApoE3, ApoE4 or ApoE3^Ch^ particles, analyzed for ethidium homodimer-1 (EthD-1) intensity and normalized to NaOH (Aβ) or DMSO (erastin) treated control neurons without particles (dashed line). n = 4 independent experiments; mean ± SEM. One-way ANOVA with Tukey’s post-test. For all graphs independent replicates are in large shapes and technical replicates in small shapes; *p < 0.05, **p < 0.01, ***p < 0.001, ****p < 0.0001.

We next turned to a more direct assay to measure endogenous lipid hydroperoxides. Even in the absence of exogenous particles, ApoE4 neurons from targeted replacement rats or derived from human induced pluripotent stem cells (iPSCs) had elevated levels of lipid hydroperoxides (Figures 2C and 2D). This elevation in lipid hydroperoxides was rescued with the addition of ApoE2 or ApoE3Ch particles by promoting their release into the media (Figures 2E-2J and S4A-S4D). ApoE3 particles were effective in effluxing lipid hydroperoxides from wild-type (Figures S4A-S4D) but not ApoE4 neurons (Figures 2E-2J). ApoE4 particles not only failed to extract lipid hydroperoxides but promoted lipid peroxide accumulation in all genotypes tested (Figures 2E-2H, S4A and S4B). Since soluble Aβ in the extracellular space increases in AD and correlates with disease progression^20,21^, we also tested whether ApoE particles affect lipid peroxidation in the presence of Aβ42. The addition of Aβ42 further increased the levels of lipid hydroperoxides in ApoE4 neurons, which was exacerbated with ApoE4 particles but rescued with ApoE2 particles (Figure 2K).

We then examined the effects of particles on neuronal health by staining with ethidium homodimer-1, a marker of cell death, and measuring fluorescence on a plate reader. Consistently, ApoE2 and ApoE3Ch particles rescue neurons from Aβ42 and erastin-induced death as indicated by reduced ethidium homodimer-1 fluorescence (Figures 2L-2O and S4E). Conversely, ApoE4 particles further increased neuronal death (Figures 2L, 2M and S4E). As erastin is a potent inducer of ferroptosis, this data indicates that ApoE2 and ApoE3Ch particles protect neurons from ferroptosis by lipid hydroperoxide efflux, while ApoE4 particles promote ferroptosis by exacerbating lipid hydroperoxide accumulation.

### ApoE2 and ApoE3Ch extract peroxidated lipids via ABCA7 transporters

We next wondered how ApoE2 and ApoE3Ch particles mediate peroxidated lipid release. Since these protective variants are known to have reduced receptor binding, we reasoned they could be acting though adenosine triphosphate-binding cassette (ABC) transporters on the cell surface that are known to efflux lipids onto lipoprotein particles. Two of these transporters, ABCA1 and ABCA7, are expressed by neurons, contribute to neuron-to-glia lipid transport, and have been identified as a risk factor for Alzheimer’s disease^22^, but whether they expel peroxidated lipids is unknown.

To explore the role of these transporters in lipid efflux, we knocked down ABCA1 or ABCA7 in neurons using lentivirus to deliver three independent shRNAmiRs for each target protein or a non-targeting shRNAmiR control (Figures 3A-3D). We discovered a robust reduction in ApoE2 and ApoE3Ch-mediated efflux of unsaturated fatty acids with ABCA7 knockdown (Figures 3E and 3F). Similarly, ABCA7 knockdown prevented ApoE2 and ApoE3Ch-mediated peroxidated lipid efflux resulting in their accumulation in neurons (Figures 3G-3J). There was no effect of ABCA7 knockdown on the release of unsaturated fatty acids in the presence of ApoE4 particles (Figure S5A). ABCA1 knockdown also had no effect on unsaturated fatty acid release (Figures S5B and S5C), lipid hydroperoxide release or accumulation in neurons with the addition of ApoE2 or ApoE4 particles (Figures 3K, 3L, S5D and S5E).

**Figure 3.**
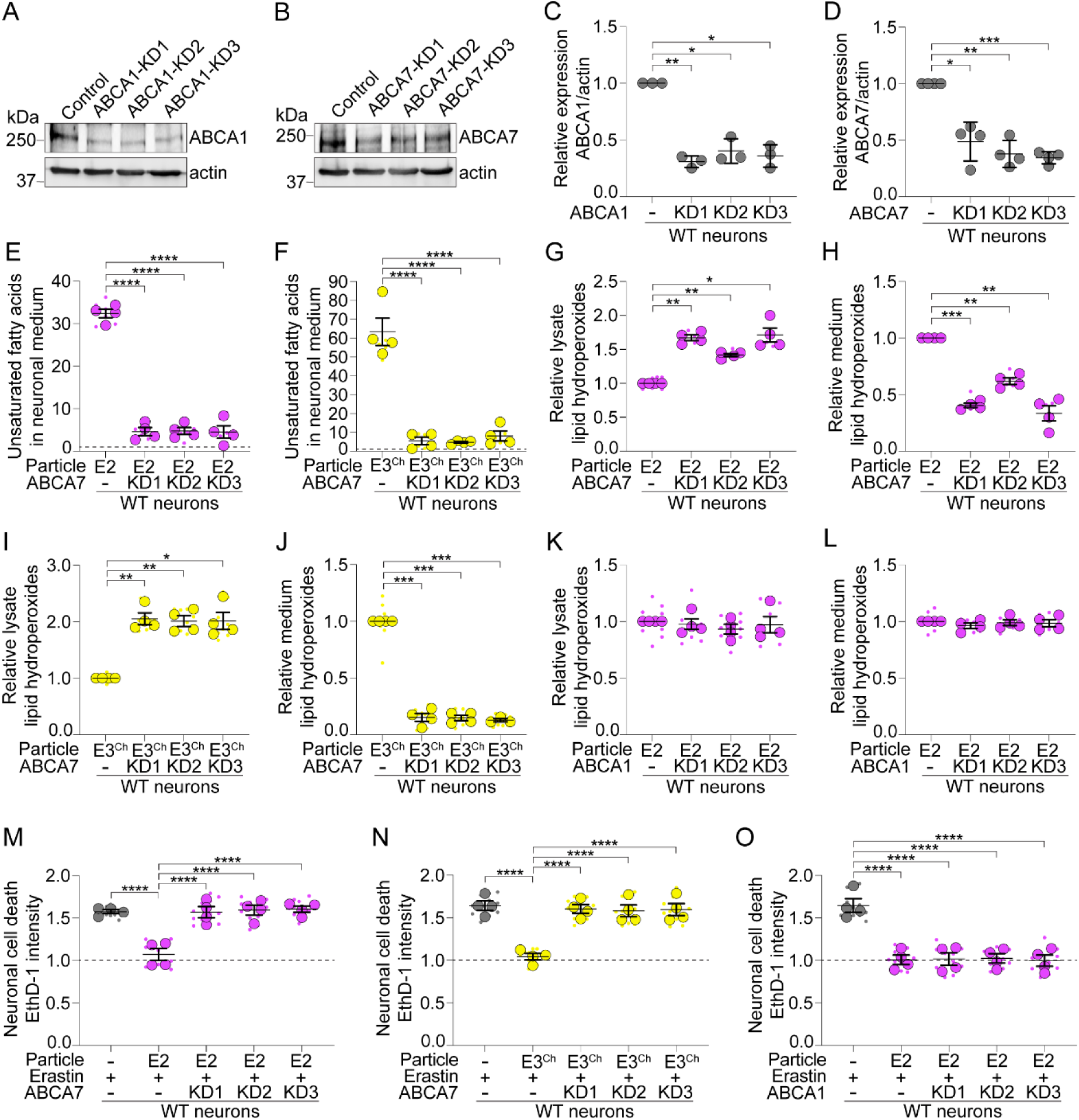
ApoE2 and ApoE3Ch particles extract peroxidated lipids through ABCA7 (A) WT neurons were transduced with lentivirus expressing non-targeting shRNAmiR (control) or ABCA1 targeting shRNAmiRs (KD1-3). Cell lysates were analyzed by Western blot for ABCA1 levels and β-actin. (B) WT neurons were transduced with lentivirus expressing non-targeting shRNAmiR (control) or ABCA7 targeting shRNAmiRs (KD1-3). Cell lysates were analyzed by Western blot for ABCA7 levels and β-actin. (C) Levels of ABCA1/β-actin were normalized to non-targeting control. n = 3 independent experiments; mean ± SD. One sample t test with Bonferroni correction. (D) Levels of ABCA7/β-actin were normalized to non-targeting control. n = 4 independent experiments; mean ± SD. One sample t test with Bonferroni correction. (E and F) WT neurons treated with ABCA7 or non-targeting shRNAmiR. After ApoE2 or ApoE3^Ch^ particle treatment, conditioned HBSS was analyzed for endogenous unsaturated fatty acids and normalized to non-targeting shRNAmiR control neurons without particles (dashed line). n = 4 independent experiments; mean ± SEM. One-way ANOVA with Tukey’s post-test. (G and H) WT neurons treated with ABCA7 or non-targeting shRNAmiR. After ApoE2 treatment, lysates (G) and conditioned HBSS (H) were analyzed for lipid hydroperoxides and normalized to non-targeting shRNAmiR control neurons treated with ApoE2 particles. n = 4 independent experiments; mean ± SEM. One sample t test with Bonferroni correction. (I and J) WT neurons treated with ABCA7 or non-targeting shRNAmiR. After ApoE3^Ch^ treatment, lysates (I) and conditioned HBSS (J) were analyzed for lipid hydroperoxides and normalized to non-targeting shRNAmiR control neurons treated with ApoE3^Ch^ particles. n = 3-4 independent experiments; mean ± SEM. One sample t test with Bonferroni correction. (K and L) WT neurons treated with ABCA1 or non-targeting shRNAmiR. After ApoE2 treatment, lysates (K) and conditioned HBSS (L) were analyzed for lipid hydroperoxides and normalized to non-targeting shRNAmiR control neurons treated with ApoE2 particles. n = 4 independent experiments; mean ± SEM. One sample t test with Bonferroni correction. (M and N) WT neurons treated with ABCA7 or non-targeting shRNAmiR. After erastin with or without ApoE2 or ApoE3^Ch^ particle treatment in media, cells were analyzed for ethidium homodimer-1 (EthD-1) intensity and normalized to non-targeting shRNAmiR DMSO treated control neurons (dashed line). n = 4 independent experiments; mean ± SEM. One-way ANOVA with Tukey’s post-test. (O) WT neurons treated with ABCA1 or non-targeting shRNAmiR. After erastin with or without ApoE2 particle treatment in media, cells were analyzed for ethidium homodimer-1 (EthD-1) intensity and normalized to non-targeting shRNAmiR DMSO treated control neurons (dashed line). n = 4 independent experiments; mean ± SEM. One-way ANOVA with Tukey’s post-test. For all graphs independent replicates are in large shapes and technical replicates in small shapes; *p < 0.05, **p < 0.01, ***p < 0.001, ****p < 0.0001.

We next wondered if ABCA7 prevents cell death downstream of lipid peroxidation. While ABCA7 knockdown alone had no effect on neuronal cell death (Figure S5F), loss of ABCA7 prevented ApoE2 and ApoE3Ch from protecting neurons from ferroptosis in the presence of erastin (Figures 3M and 3N). Conversely, ABCA1 knockdown had no effect on ApoE2-mediated neuron survival in the presence of erastin (Figure 3O). Collectively, these data reveal that ApoE2 and ApoE3Ch particles exert their protective actions by selectively extracting peroxidated lipids through the ABCA7 transporter.

### ApoE4-mediated endolysosomal dysfunction rescued by protective particles

Since ApoE4 particles exacerbate peroxidated lipid accumulation^16^, that led us to ask whether this might occur by dysregulation of the endolysosomal system^23^. For this to occur, particles must first be endocytosed. While ApoE isoforms can have varying affinities for some receptors^24–26^, neurons internalize equivalent amounts of ApoE particles regardless of the isoform (Figures S6A and S6B). We then measured lipid hydroperoxide levels in the presence of the endocytosis inhibitor Pitstop 2. Indeed, Pitstop 2 prevented the increase in lipid hydroperoxide accumulation caused by ApoE4 particles treated with or without erastin (Figures 4A, S6C and S6D). Blocking endocytosis with Pitstop 2 had no effect on lipid hydroperoxide levels in neurons treated with ApoE2 particles (Figures 4B, S6E and S6F). This indicates that endocytosis is required for the toxic effects of ApoE4 particles on neurons. On the other hand, the protective actions of ApoE2 particles on reducing neuronal lipid peroxidation are independent of endocytosis, consistent with acting on transporters on the cell surface.

**Figure 4.**
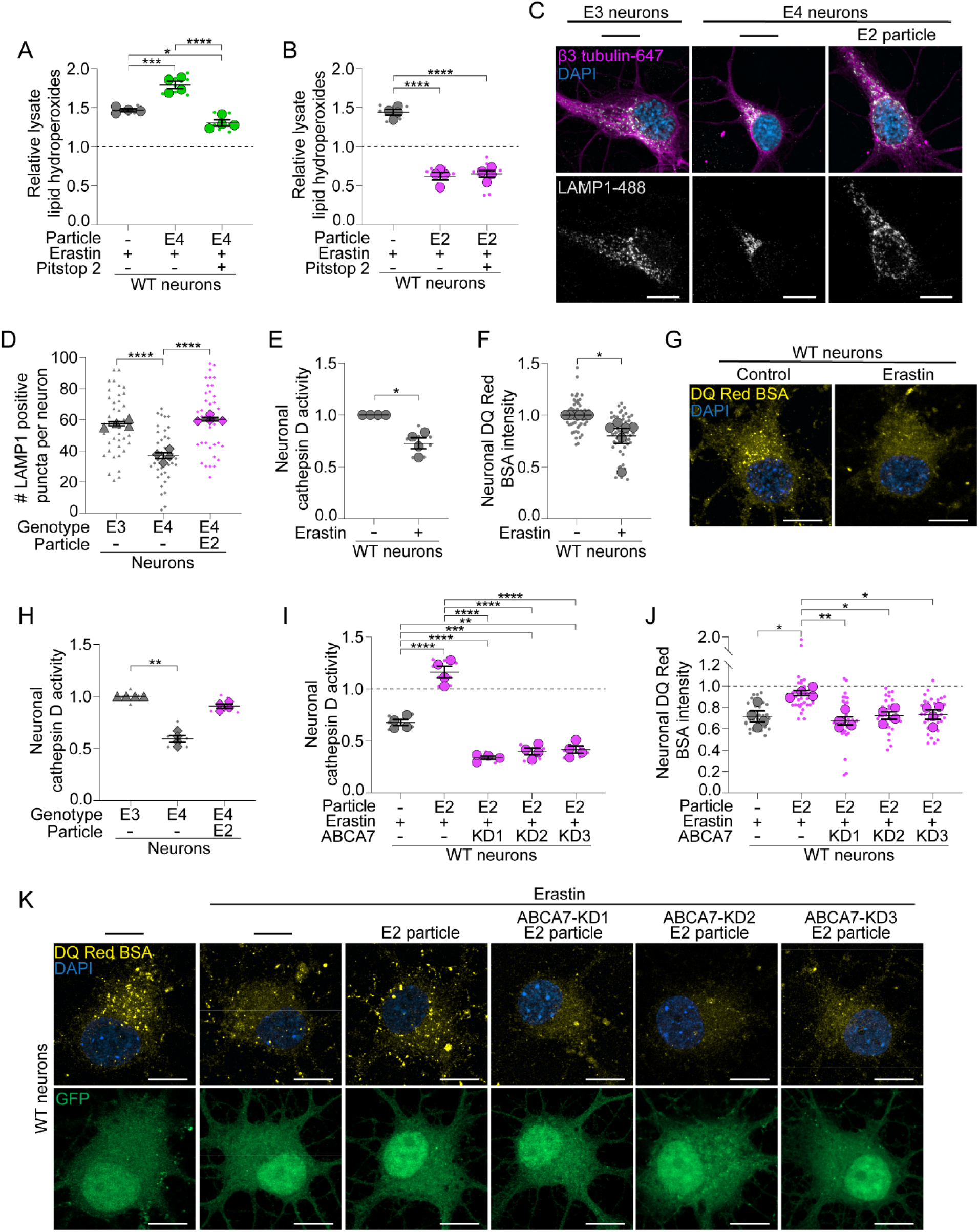
ApoE2 and ApoE3Ch rescue ApoE4-mediated endolysosomal dysfunction (A and B) WT neurons after treatment in media with ApoE2 or ApoE4 particles with or without Pitstop 2 and erastin. Lysates analyzed for lipid hydroperoxides and normalized to DMSO treated control neurons (dashed line). n = 4 independent experiments; mean ± SEM. One-way ANOVA with Tukey’s post-test. (C and D) Airyscan maximum intensity projection images of ApoE3 or ApoE4 neurons with or without ApoE2 particles in HBSS, immunostained for LAMP1 and the number of LAMP1 positive puncta were quantified. n = 4 independent experiments; mean ± SEM. One-way ANOVA with Tukey’s post-test. Scale bars are 10 μm. (E) WT neurons treated with erastin in HBSS were analyzed for cathepsin D activity and normalized to DMSO treated control neurons. n = 4 independent experiments; mean ± SEM. One sample t test with Bonferroni correction. (F) WT neurons treated with erastin in HBSS were analyzed for DQ Red BSA intensity and normalized to DMSO treated control neurons. n = 6 independent experiments; mean ± SEM. One sample t test with Bonferroni correction. (G) Airyscan images of WT neurons displayed as maximum intensity projections following treatment in HBSS with or without erastin, labeled with DQ Red BSA. Scale bars are 10 μm. (H) ApoE3 and ApoE4 neurons with or without ApoE2 particles in HBSS analyzed for cathepsin D activity and normalized to ApoE3 neurons. n = 4 independent experiments; mean ± SEM. One sample t test with Bonferroni correction. (I) WT neurons treated with ABCA7 or non-targeting shRNAmiR. After erastin treatment in media with or without ApoE2 particles, cells were analyzed for cathepsin D activity and normalized to non-targeting shRNAmiR DMSO treated control neurons (dashed line). n = 4 independent experiments; mean ± SEM. One-way ANOVA with Tukey’s post-test. (J) WT neurons treated with ABCA7 or non-targeting shRNAmiR. After erastin treatment in media with or without ApoE2 particles, cells were analyzed for DQ Red BSA intensity and normalized to non-targeting shRNAmiR DMSO treated control neurons (dashed line). n = 4 independent experiments; mean ± SEM. One-way ANOVA with Tukey’s post-test. (K) Airyscan images of WT neurons treated with ABCA7 or non-targeting shRNAmiR (GFP) displayed as maximum intensity projections following treatment in media with or without erastin or ApoE2 particles, labeled with DQ Red BSA. Scale bars are 10 μm. For all graphs independent replicates are in large shapes and technical replicates in small shapes; *p < 0.05, **p < 0.01, ***p < 0.001, ****p < 0.0001.

Since ApoE4 particles exert their deleterious action intracellularly, this raised the possibility that ApoE4 may disrupt alternate pathways for lipid efflux, such as lysosomal exocytosis^27^. However, ApoE4 particles had no effect on the release of iron (Figures S6G and S6H) or Red-C12 (Figures 1A and S3A-S3C), which occurs during lysosomal exocytosis^27^, indicating that ApoE4 is unlikely to be affecting this pathway. Instead, ApoE4 could disrupt endolysosomal physiology through its effects on lipid peroxidation. For example, erastin can reduce the number of functional lysosomes^28^. We observed a similar decrease in LAMP1-positive lysosomes in ApoE4 neurons or wild-type neurons treated with ApoE4 particles (Figures 4C, 4D, S6I and S6J). Conversely, ApoE2 particles increased the number of LAMP1-positive lysosomes (Figures 4C, 4D, S6I and S6J). Consistent with fewer lysosomes, lipid peroxidation induced by erastin decreased the degradative capacity of the neurons, as indicated by decreased activity of the lysosomal protease cathepsin D (Figure 4E) and a reduction in DQ Red BSA fluorescence, a lysosomal protease substrate that fluoresces upon proteolytic cleavage in lysosomes (Figures 4F and 4G). This indicates that increased lipid peroxidation diminishes lysosomal function. Consistent with exacerbating lipid peroxidation, cathepsin D activity and DQ Red BSA intensity was also reduced in ApoE4 neurons or in wild-type neurons treated with ApoE4 particles (Figures 4H and S6K-S6M). In contrast, ApoE2 particles restore cathepsin D activity and DQ Red BSA intensity in ApoE4 neurons and wild-type neurons (Figures 4H and S6K-S6M), which was dependent on ABCA7 transporters (Figures 4I-4K). Collectively, this data indicates that ApoE4 particles exert their damaging effects by disrupting the endolysosomal system which can be restored by ApoE2 particles acting on the surface.

### ApoE2 and ApoE3Ch particles protect intact hippocampal tissue from lipid peroxidation

Since ApoE2 and ApoE3Ch particles rescue cultured neurons from toxicity associated with increased lipid peroxidation, we next sought to test whether they would be similarly effective at protecting neural tissue. We used intact hippocampi from young, targeted replacement rats as our model system, as they continue to exhibit neuronal activity ex vivo for several hours^29,30^. To stimulate neural activity, we perfused the hippocampi with oxygenated artificial cerebrospinal fluid containing low magnesium and high potassium (Figure 5A). Consistent with our in vitro results, perfused hippocampi from ApoE4 animals have elevated lipid hydroperoxides and reduced cathepsin D activity (Figures 5B and 5C).

**Figure 5.**
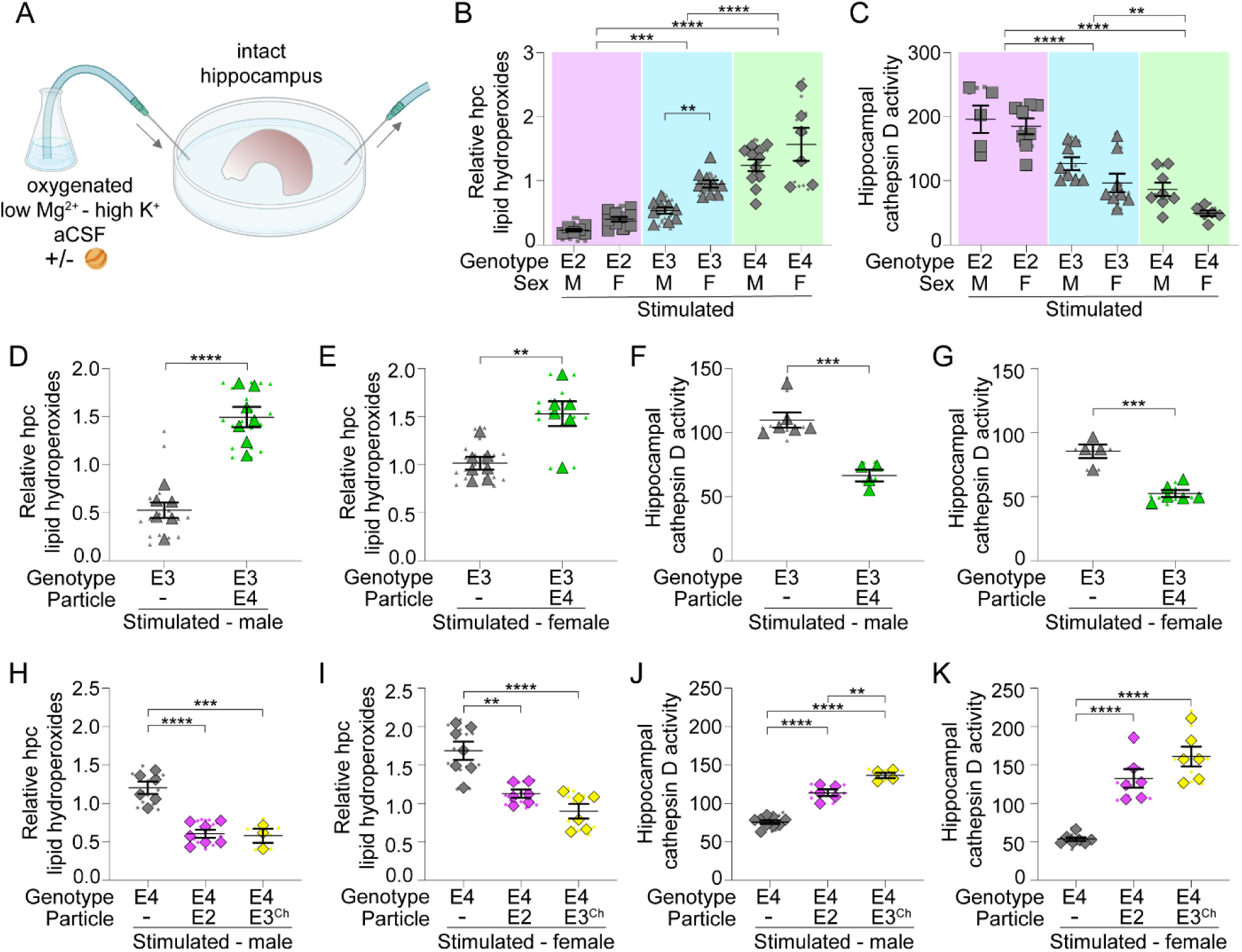
ApoE2 and ApoE3Ch particles rescue hippocampi from lipid-mediated toxicity (A) Schematic of the intact hippocampal experiment. (B) Intact hippocampi from ApoE targeted replacement rats perfused with low Mg^2+^-high K^+^ artificial cerebrospinal fluid (aCSF) analyzed for lipid hydroperoxides and normalized to tissue weight. Minimum n = 6 animals per group; mean ± SEM. Two-way ANOVA with Tukey’s post-test for genotype comparisons and Šidák’s post-test for sex-specific comparisons. M, male; F, female. Hpc, hippocampal. (C) Intact hippocampi from ApoE targeted replacement rats perfused with low Mg^2+^ - high K^+^ aCSF analyzed for cathepsin D activity and normalized to protein content. Minimum n = 5 animals per group; mean ± SEM. Two-way ANOVA with Tukey’s post-test for genotype comparisons and Šidák’s post-test for sex-specific comparisons. M, male; F, female. (D and E) Intact hippocampi from ApoE3 rats perfused with low Mg^2+^ - high K^+^ aCSF with or without ApoE4 particles, analyzed for lipid hydroperoxides and normalized to tissue weight. Minimum n = 6 animals per group; mean ± SEM. Two-tailed Student’s t test. (F and G) Intact hippocampi from ApoE3 rats perfused with low Mg^2+^ - high K^+^ artificial aCSF with or without ApoE4 particles, analyzed for cathepsin D activity and normalized to protein content. Minimum n = 4 animals per group; mean ± SEM. Two-tailed Student’s t test (H and I) Intact hippocampi from ApoE4 rats perfused with low Mg^2+^ - high K^+^ aCSF treated with or without ApoE2 or ApoE3^Ch^ particles, analyzed for lipid hydroperoxides and normalized to tissue weight. Minimum n = 3 animals per group; mean ± SEM. One-way ANOVA with Tukey’s post-test. (J and K) Intact hippocampi from ApoE4 rats perfused with low Mg^2+^ - high K^+^ aCSF treated with or without ApoE2 or ApoE3^Ch^ particles, analyzed for cathepsin D activity and normalized to protein content. Minimum n = 4 animals per group; mean ± SEM. One-way ANOVA with Tukey’s post-test. For all graphs independent replicates are in large shapes and technical replicates in small shapes; *p < 0.05, **p < 0.01, ***p < 0.001, ****p < 0.0001.

We next explored how ApoE particles affect the stimulated hippocampal tissue. ApoE4 particles induced dysfunction in ApoE3 tissue as evidenced by increased lipid hydroperoxides and decreased cathepsin D activity, regardless of the sex (Figures 5D-5G). ApoE2 and ApoE3Ch, conversely, rescued ApoE4 tissue from dysfunction by lowering lipid hydroperoxides and increasing cathepsin D activity in both male and female tissue (Figures 5H-5K). Together, these results support the role of ApoE2 and ApoE3Ch in protecting intact hippocampal tissue from the damaging effects of lipid peroxidation.

### ApoE2 and ApoE3Ch particles restore hippocampal activity

Since ApoE2 and ApoE3Ch particles prevent lipid peroxidation and endolysosomal dysfunction in intact hippocampi, we next sought to determine if they also support neuronal function. To this end, we assessed the effects of ApoE on neural activity by measuring local field potentials in the subiculum following stimulation with artificial cerebrospinal fluid containing low magnesium and high potassium (Figures 6A and 6B). The subiculum was chosen as it is one of the earliest regions within the hippocampus to be affected in AD^31^. Both ApoE3 and ApoE4 hippocampi increased in firing upon stimulation (Figures 6C-6F). Consistent with previous studies^32^, we then observed a sharp drop in ApoE4 activity as this tissue was unable to sustain stimulated activity as compared to ApoE3 (Figures 6C-6F). Both the frequency and amplitude of firing events was reduced in ApoE4 tissue as compared to ApoE3 one hour after stimulation (Figures 6C and 6G-6J). We next examined whether we could restore the stimulated activity of the ApoE4 tissue with the addition of the protective particles. Indeed, both ApoE2 and ApoE3Ch particles recovered the frequency and amplitude of firing in stimulated ApoE4 tissue (Figures 6C-6J). Collectively, these data reveal that ApoE4 promotes dysfunctional lipid handling in the intact hippocampus which results in impaired ability to sustain neuronal activity. Protective ApoE2 and ApoE3Ch particles rescue the ApoE4-induced lipid defects and thereby recover deficits in stimulated neural activity.

**Figure 6.**
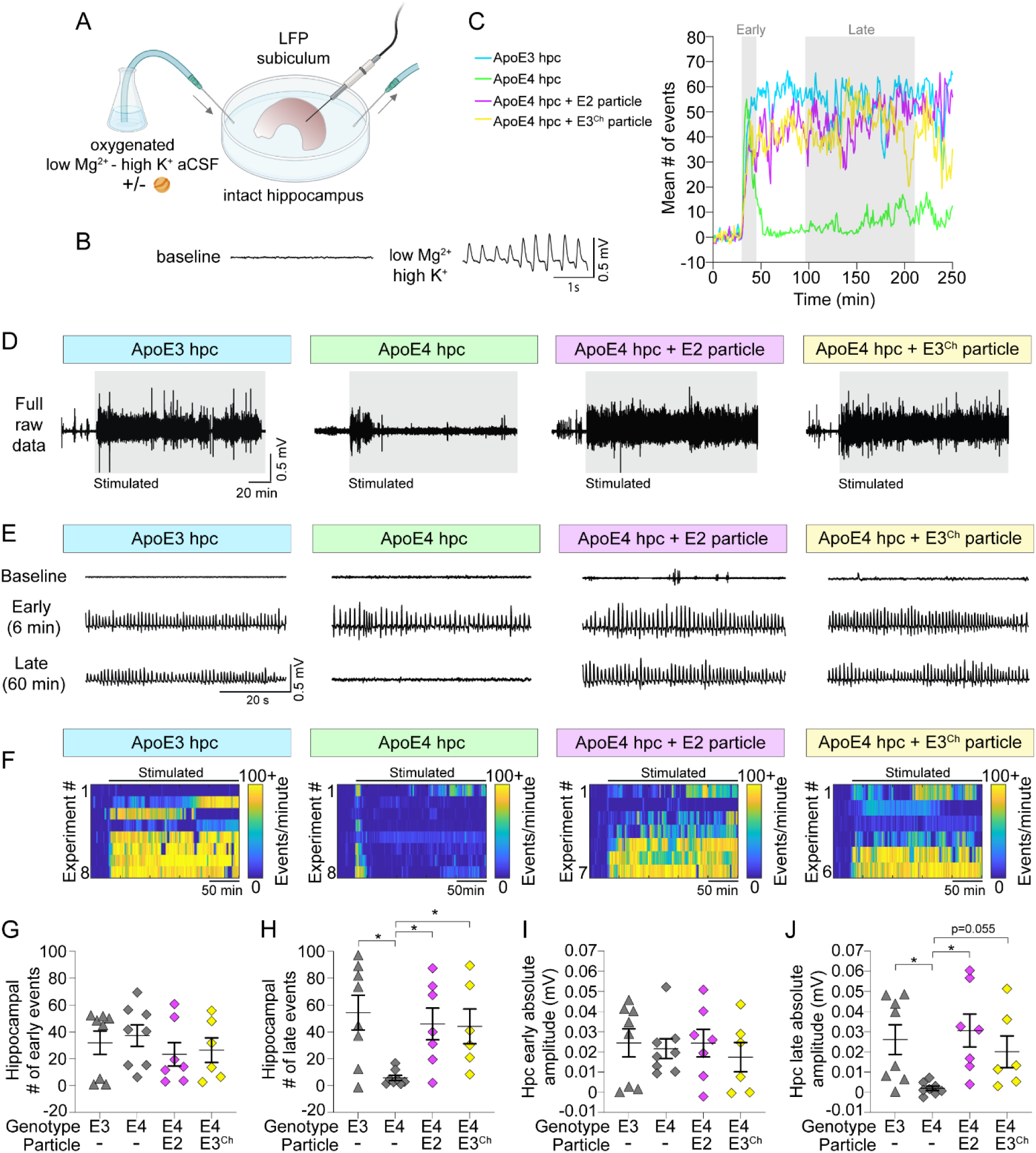
ApoE2 andApoE3Ch particles rescue impaired hippocampal activity (A) Schematic of intact hippocampal electrophysiology recordings. (B) Representative raw LFP data of intact hippocampi from ApoE3 rat during baseline and perfusion with low Mg^2+^ - high K^+^ artificial cerebrospinal fluid (aCSF). (C) Frequency of events of intact hippocampi from ApoE3 or ApoE4 rats during baseline and perfusion with low Mg^2+^ - high K^+^ aCSF with or without ApoE2 or ApoE3^Ch^ particles. Hpc, hippocampal. (D) Representative raw LFP data of intact hippocampi from ApoE3 or ApoE4 rats during baseline and perfusion with low Mg^2+^ - high K^+^ aCSF with or without ApoE2 or ApoE3^Ch^ particles. (E) Expanded representative LFPs during baseline (top), early (middle) and late (bottom) perfusion with low Mg^2+^ - high K^+^ aCSF with or without ApoE2 or ApoE3^Ch^ particles. (F) Frequency of events per minute for each experiment of intact hippocampi from ApoE3 or ApoE4 rats during baseline and perfusion with low Mg^2+^ - high K^+^ aCSF with or without ApoE2 or ApoE3^Ch^ particles. n = 6-8 hippocampi. (G and H) Intact hippocampi from ApoE3 or ApoE4 rats perfused with low Mg^2+^ - high K^+^ aCSF with or without ApoE2 or ApoE3^Ch^ particles, analyzed for frequency of events early (G) or late (H). Minimum n = 6 hippocampi per group; mean ± SEM. Kruskal-Wallis with Dunn’s post-test. (I and J) Intact hippocampi from ApoE3 or ApoE4 rats perfused with low Mg^2+^ - high K^+^ aCSF with or without ApoE2 or ApoE3^Ch^ particles, analyzed for amplitude of events early (I) or late (J). Minimum n = 6 hippocampi per group; mean ± SEM. Kruskal-Wallis with Dunn’s post-test.

## DISCUSSION

Neuron to glia lipid transport is critical for protecting the brain from lipid-mediated toxicity. Here, we uncovered a key mechanism by which ApoE isoforms as a component of extracellular lipoprotein particles interact with neurons to facilitate this process. We discovered that the AD-protective ApoE2 and ApoE3Ch isoforms protect neurons from toxicity by facilitating efflux of peroxidated unsaturated fatty acids through the ABCA7 transporter on the plasma membrane^33^. These protective particles can rescue ApoE4 neurons from elevated lipid hydroperoxides, impaired endolysosomal function and cell death by ferroptosis.

Lipid peroxidation is an early pathological hallmark of AD^12^. Here, we discovered a critical function for ApoE2 and ApoE3Ch particles: efflux of peroxidated unsaturated lipids from both human and rat neurons. During oxidative stress, lipids become peroxidated in the presence of iron and excess reactive oxygen species driving ferroptosis. While polyunsaturated phospholipids such as phosphatidylethanolamine are the preferred substrate for peroxidation^12^, the species of unsaturated lipids being peroxidated and extracted by ApoE particles from neurons remains to be determined. What is clear is that exogenous ApoE2 and ApoE3Ch particles are potent inhibitors of neuronal ferroptotic cell death via their peroxidated lipid efflux function.

Genome-wide association studies have identified lipid metabolism genes as major risk factors for late onset AD^34,35^. Like ApoE4, several of these genes are similarly involved in the transport of lipids from neurons to their destination in glial lipid droplets^22^. Notably, the ABC transporters, ABCA1 and ABCA7 when expressed by neurons participate in glial lipid droplet formation^22^. These transporters act on the cell surface and transfer lipids to the outer leaflet of membranes where they can be exported onto extracellular ApoE particles. Here, we show that ABCA7 is required for the efflux of peroxidated unsaturated fatty acids from neurons by ApoE2 and ApoE3Ch particles. This is consistent with work showing that ABCA7 is more effective at effluxing phospholipids as opposed to cholesterol^36^. What remains unclear is how peroxidated lipids are the preferred substrate for this transport, and whether the increased affinity for peroxidated acyl chains is mediated by ABCA7, ApoE2 or ApoE3Ch.

We also found that ABCA1 did not affect peroxidated lipid efflux or ferroptotic cell death. This was surprising given that loss of the ABCA1 homolog Eato in *Drosophila* results in loss of glial lipid droplets and neurodegeneration which can be rescued with an antioxidant^22^. Eato may be better able to efflux peroxidated phospholipids in *Drosophila* compared to ABCA1 in mammals. Alternatively, there may be differences in ABCA1 trafficking in different model systems. For example, in ApoE4 rodent cells ABCA1 becomes trapped in late endosomes and is no longer recycled to the plasma membrane to mediate lipid efflux^37^.

We also found that protective particles rescue ApoE4-dependent impairment of the endolysosomal pathway. Endolysosomal dysfunction contributes to AD pathogenesis^38,39^ and can be caused by ApoE4 genotype independent of Aβ pathology^40^. We show that endocytosis of exogenous ApoE4 particles is sufficient to reduce the number of functional lysosomes and their degradative capacity through its effects on lipid peroxidation. Interestingly, all isoforms of ApoE were readily taken up by neurons, but only ApoE4 induced lysosomal dysfunction. Protective particles rescue ApoE4-induced endolysosomal dysfunction through ABCA7-dependent lipid efflux. Since the protective particles are acting on ABCA7 on the cell surface, this could reduce the amount of peroxidated lipids trafficked to the lysosomes from endosomes originating at the plasma membrane, as well as via autophagy. In addition to its role in the endolysosomes, ABCA7 also affects mitochondrial health. Specifically, loss of ABCA7 in iPSC-derived neurons decreased mitochondria-related phospholipids and altered mitochondria morphology, resulting in impaired mitochondrial respiration and excess reactive oxygen species production^41^. These mitochondrial defects in turn can have profound consequences on neuronal function as iPSC-derived neurons had decreased spontaneous synaptic activity^41^. This is consistent with our electrophysiology data showing a similar decrease in neuronal activity with ApoE4 that can be rescued with the protective particles in intact hippocampal tissue.

## ACKNOWLEDGMENTS

Experiments were performed at the University of Alberta Faculty of Medicine & Dentistry Cell Imaging Core, RRID:SCR_019200. iPSC work was performed at the Center for Neurogenomics and Cognitive Research, Vrije Universiteit Amsterdam. We thank Luis Rubio-Atonal for assistance with ex vivo assays. I.R. is supported by a Canada Graduate Scholarship from the Canadian Institutes of Health Research Doctoral Award #181551 and an Izaak Walton Killam Memorial Scholarship. F.M.F is supported by a ZonMW memorabel fellowship (733050515) and Alzheimer Nederland pilot grant (WE.03-2019-13). R.V.D.K is supported by the Chan Zuckerberg Initiative, collaborative pairs grant DAF2022-250616 and a Vidi 2021 (09150172110086) from the Dutch Research Council (NWO). M.S.I. is supported by a sponsored research contract from Kisbee Therapeutics, and grants from the Canadian Institutes of Health Research (grant#173321) and the Women and Children’s Health Research Institute.

## AUTHOR CONTRIBUTIONS

Conceptualization, I.R. and M.S.I.; Methodology, I.R., W.C., K.C., C.G., J.J., E.R., and M.S.I.; Investigation, I.R., A.D.D., J.B., F.M.F., W.C., J.C., and C.G.; Analysis, I.R., A.D.D., W.C., and J.J.; Writing – Original Draft, I.R. and M.S.I.; Writing – Review & Editing, I.R., A.D.D., J.B., F.M.F., W.C., J.C., K.C., J.J., E.R., and M.S.I.; Supervision, R.V.D.K., J.J., E.R., and M.S.I.

## DECLARATION OF INTERESTS

Partial funding for this project comes from Kisbee Therapeutics, where E.R. and C.G. are employees and hold a patent related to the therapeutic use of ApoE particles.

## STAR METHODS

### KEY RESOURCES TABLE

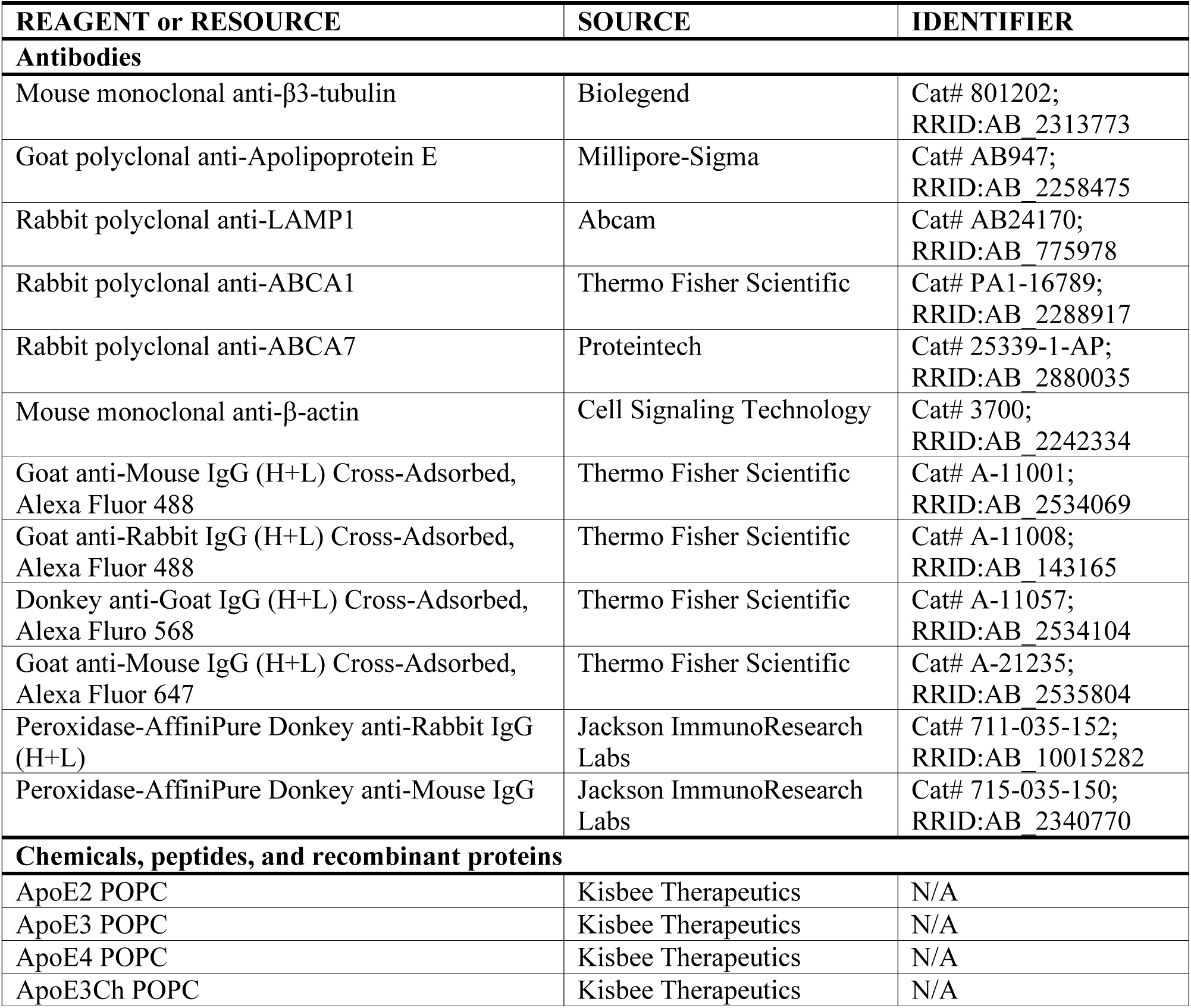
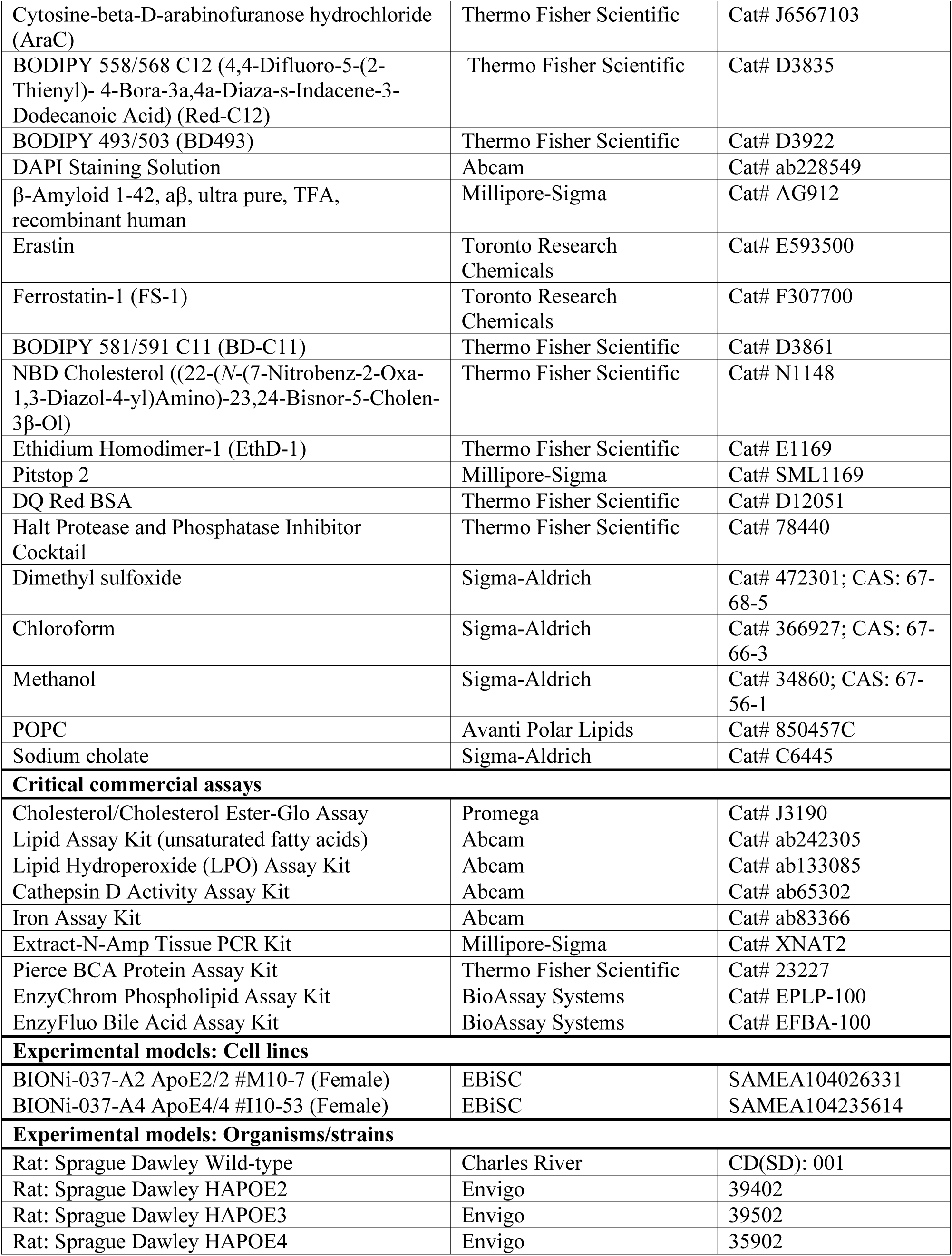
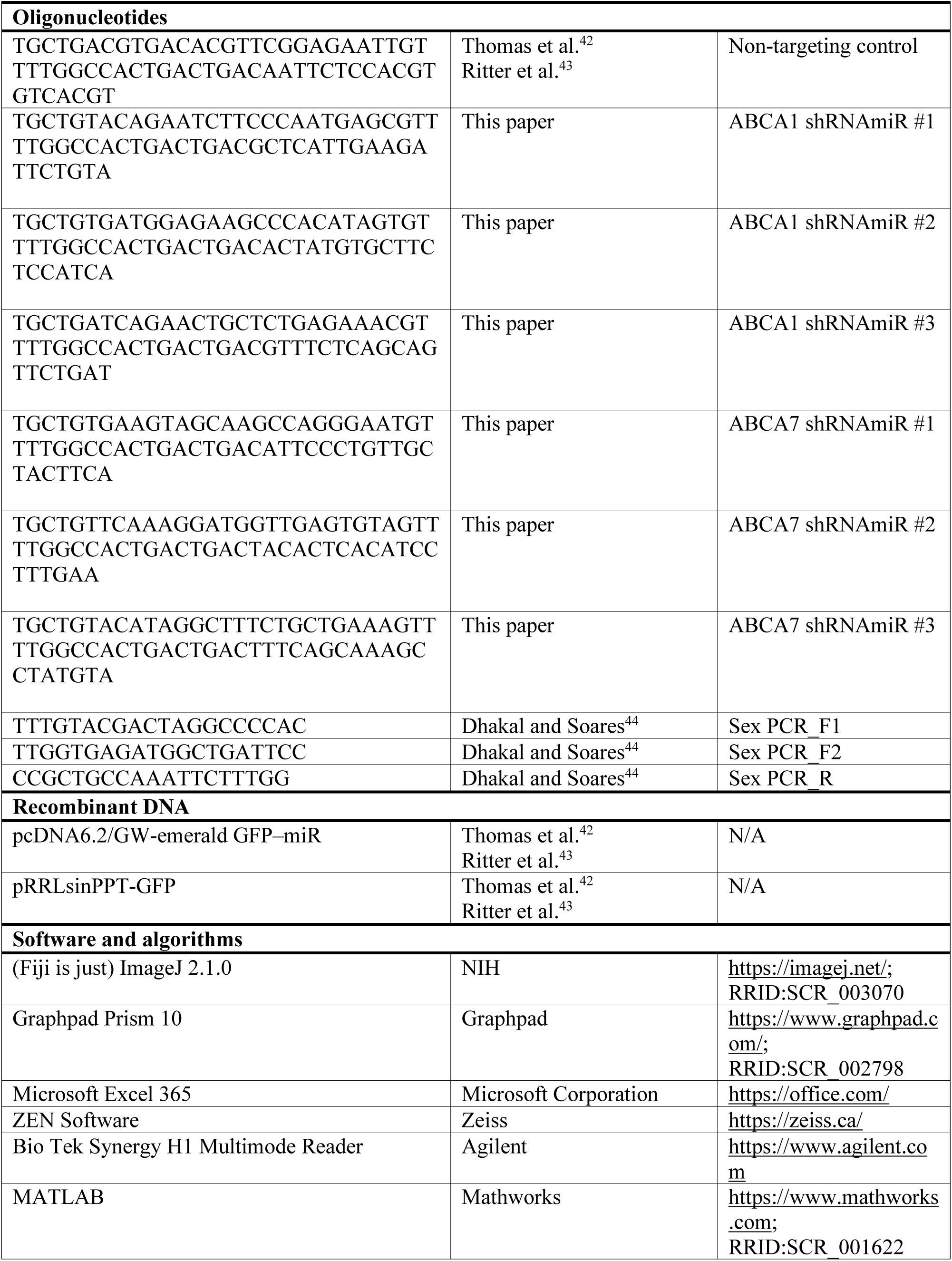
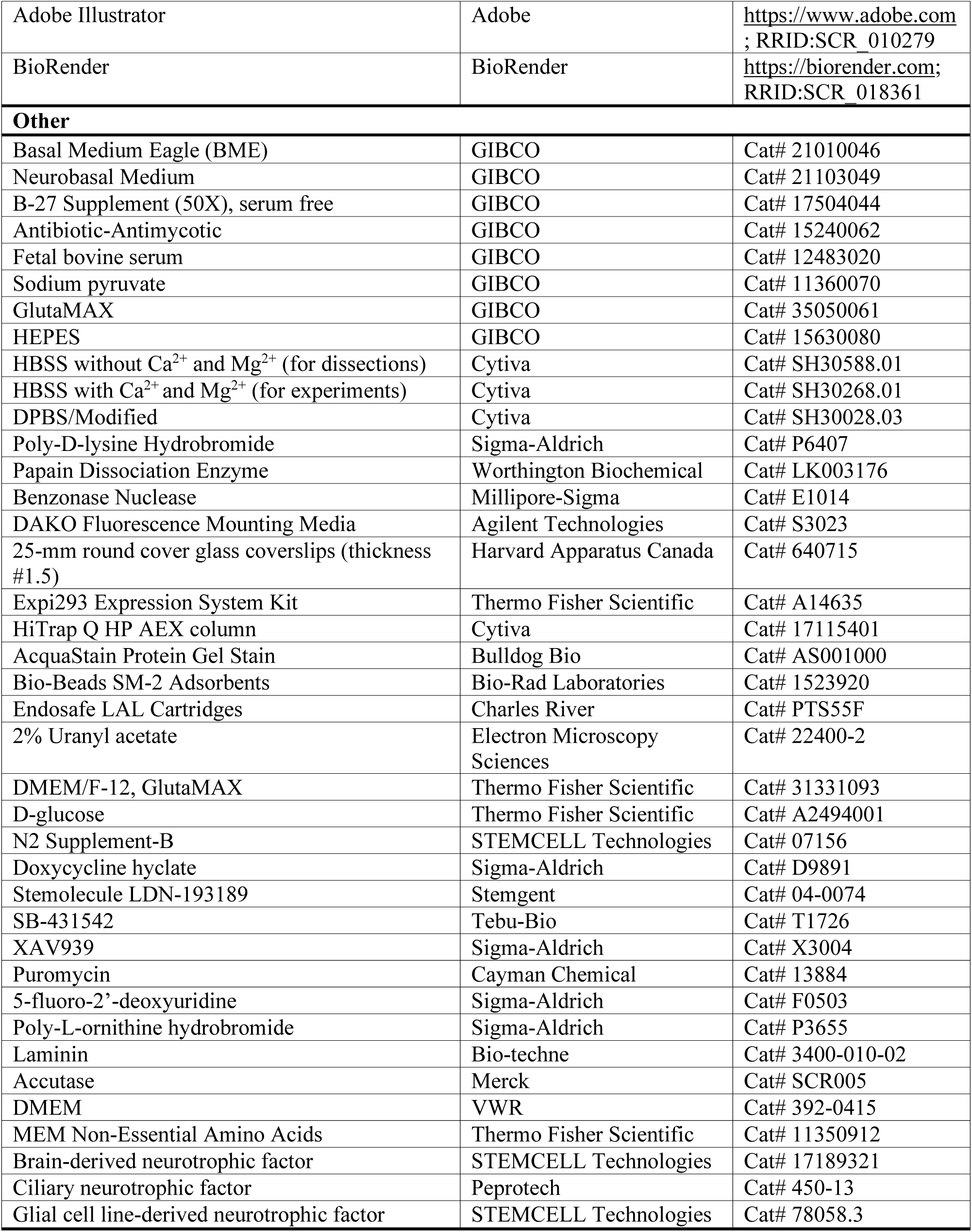

### RESOURCE AVAILABILITY

#### Lead contact

Further information and requests for resources and reagents should be directed to and will be fulfilled by the lead contact, Maria S. Ioannou.

#### Materials availability

This study did not generate new unique reagents.

#### Data and code availability

- All data reported in this paper will be shared by the lead contact upon request.
- This paper does not report original code.
- Any additional information required to reanalyze the data reported in this paper is available from the lead contact upon request.

## EXPERIMENTAL MODEL AND SUBJECT DETAILS

### Animals

All animal work was approved by and performed in accordance with the Canadian Council of Animal Care at the University of Alberta (AUP#3358). Sprague-Dawley timed pregnant rats were obtained from Charles River Laboratories and arrived at our facility one week prior to birth. Targeted replacement human ApoE2, ApoE3 and ApoE4 rats were bred in house. P0-P1 pups were used for primary neuron and glia cultures. Ex vivo electrophysiology experiments were performed on P9-P10 targeted replacement human ApoE rats. All experiments were performed on male and female animals. No restrictions were imposed on food and water.

### Primary culture of hippocampal neurons and glia

Primary hippocampal cultures were prepared as previously described (Ioannou et al.^45^). Briefly, hippocampi were dissected from P0-P1 rat pups and digested with papain and benzonase nuclease. After digestion, the tissues were gently triturated and filtered with a 70-µm nylon cell strainer. Neurons and glia were grown on poly-D-lysine (PDL) coated 25-mm round glass coverslips for microscopy experiments or plastic tissue culture dishes for biochemical analysis. Neurons were grown in Neurobasal medium containing B-27 supplement, 2 mM GlutaMax and antibiotic-antimycotic. 1 μM AraC was added to the media 2 days after plating and maintained for 4 days to prevent growth of glia in neuronal cultures. Glia were grown in Basal Eagle Media containing 10% fetal bovine serum, 0.45% glucose, 1 mM sodium pyruvate, 2 mM GlutaMax and antibiotic-antimycotic. All cells were grown at 37°C in 5% CO_2_ and used at DIV 6-9.

### Generation of hiPSC-neurons

NGN2 transcription-based iPSC differentiation to neurons was performed as previously described (Zhang et al.^46^). iPSCs were infected in suspension (in E8 + RI) with ultra-high titer lentiviral particles provided by ALSTEM, encoding pTet-O-Ngn2-puro (Addgene #52047) and FUΔGW-rtTa (Addgene #19780). To begin neuronal induction 1 x 10^5^ cells/cm^2^ infected iPSCs were plated in N2-supplemented medium (DMEM/F-12 + GlutaMAX, 3 g/L D-glucose, 1% N2 supplement-B and 0.1% P/S) with 5 µM RI, 2 μg/ml doxycycline hyclate and dual SMAD inhibitors (100 nM LDN-193189, 10 μM SB-431542, 2 μM XAV939). On day 2, 100% of the medium was refreshed (including all day 1 supplements except RI), and 3 µg/mL puromycin was added. On day 3, 100% medium was exchanged for N2-supplemented medium with doxycycline hyclate, puromycin and 10 µM 5-fluoro-2’-deoxyuridine. 6-well plastic dishes were coated with 20 µg/mL poly-L-ornithine (PLO) overnight at room temperature followed by 3 wash steps with PBS on day 4. PLO-coated wells were subsequently coated with 5 µg/mL laminin (lam) for 2-4 h at 37°C. iPSC-derived neurons (iNeurons) were washed with 1X PBS before dissociation with accutase for 5 min at 37°C. iNeurons were collected in DMEM and pelleted by centrifugation at 180 x g for 5 min. iNeurons were resuspended and plated at 5 x 10^5^ cells/well in PLO-lam coated 6-well plastic dishes in Neurobasal medium, supplemented with 200 mM GlutaMAX, 3 g/L D-glucose, 0.5% NEAA, 2% B27, 0.1% P/S, 10 ng/mL BDNF, 10 ng/mL CNTF, and 10 ng/mL GDNF. iNeurons were cultured at 37°C with 5% CO_2_ and medium was replaced with 50% fresh medium once a week. iNeurons were used for lipid peroxidation assays 6 weeks after the start of differentiation.

### ApoE expression and purification

Expi293 cells were subcultured according to the manufacturer’s protocol. Human ApoE2, ApoE3, ApoE3Ch, and ApoE4 protein sequences were codon optimized and cloned into the pcDNA3.4 vector using GeneArt gene synthesis. Cells were transiently transfected with ExpiFectamine according to the manufacturer’s protocol. Cells were pelleted by centrifugation and then discarded. Secreted ApoE2, ApoE3, ApoE3Ch, or ApoE4 in the clarified media was purified by anion exchange chromatography and eluted with 1 M NaCl. The resulting proteins were buffer exchanged into denaturing and reducing buffer (6 M guanidinium HCl, 20 mM MOPS, 50 mM TCEP, pH 7.0) and stored at −80°C until assembly into lipoproteins. Protein purity was assessed by reducing SDS-PAGE and total protein gel staining. Protein sequences were validated with mass spectrometry (data not shown). Protein concentration was determined by measuring the sample’s absorbance at 280 nm (A280) or performing a bicinchoninic acid (BCA) assay.

### Lipid preparation

A solution of 1-palmitoyl-2-oleoyl-glycero-3-phosphocholine (POPC) in chloroform was transferred to a tared glass vial, the solvent was evaporated under a stream of nitrogen gas and residual solvent was removed under reduced pressure overnight. The lipid film was then resolubilized in buffer containing 2% w/v sodium cholate. The final concentration of resolubilized phospholipids was quantified by enzyme-coupled fluorometric assay according to the manufacturer’s protocol (BioAssay Systems).

### Lipoprotein assembly

Lipoproteins were assembled as previously described (Hubin et al.^47^). Briefly, resolubilized POPC and purified ApoE in denaturing buffer were combined to achieve a final protein concentration of 0.25–3.0 mg/mL and an ApoE/POPC ratio of 1:95–1:125 mol/mol. The solutions were mixed thoroughly by pipetting up and down several times, then allowed to incubate on ice for 1 hr. The mixtures were then transferred to dialysis devices prepared according to the manufacturer’s instructions, and the denaturant and cholate detergent were removed from the reaction mixtures via dialysis against >250-fold excess PBS buffer at 4°C. After >2 hours of dialysis, the dialysate was replaced with fresh PBS buffer and a volume of polystyrene adsorbent resin equal to 0.25-2% of the volume of the dialysate, then dialysis continued at 4°C. After >22 hours of total dialysis, the sample was retrieved from the dialysis device. The reaction mixtures were then concentrated using centrifugal spin concentrators to <20% of their original volume in a refrigerated centrifuge held at 4°C. The samples were then diluted back to their original volumes in PBS and mixed via pipetting up and down several times. This process was repeated two additional times to achieve a sufficient degree of exchange into PBS. Finally, the samples were concentrated to 1–15 mg/mL concentration and filtered through a sterile syringe filter (pore size ≤0.22 µm) in a biosafety cabinet, then stored at 4°C or frozen at −80°C until used in experiments.

### Particle characterization

ApoE particles were analyzed via native PAGE using 4-12% Tris-Glycine gels followed by total protein staining. The diameters of the lipoproteins and polydispersity of the samples were measured using a plate-based Wyatt dynamic light scattering (DLS) instrument following standard protocols. Both the diameter and morphology were also characterized by transmission electron microscopy (TEM). Samples were diluted in PBS to 0.025-0.2 mg/mL protein concentration and stained with 2% uranyl acetate. Prepared grids were analyzed on a Hitachi TEM system operating at an accelerating voltage of 100 kV. Images were captured at magnifications ranging from 20,000x to 80,000x, and particle size were determined using the highest magnification images collected for each sample (60,000x-80,000x). Diameters were determined in Fiji-ImageJ using the “Analyze Particles” function after auto local thresholding with contrast setting of 40. The protein, POPC, and cholate contents of the lipoprotein samples were measured using enzyme-coupled fluorometric or colorimetric assays. Protein content of the lipoproteins was additionally calculated by measuring the absorbance of the sample at 280 nm.

### Lentivirus production

pcDNA6.2/GW-emerald GFP–miR, pRRLsinPPT and the pRRLsinPPT-non-targeting-miR control were generous gifts from Dr. Peter McPherson, McGill University, Montreal, Quebec, Canada. Target sequences for rat ABCA1 and ABCA7 were designed using the BLOCK-iT RNAi Designer (Invitrogen). Target sequences were subcloned first into the pcDNA6.2/GW-emerald GFP–miR cassette and then into the pRRLsinPPT viral expression vector (Ritter et al.^43^). ABCA1 and ABCA7-targeting lentiviral particles were generated by VectorBuilder.

### Lentivirus transduction

Neurons were transduced with lentivirus (MOI 7.5) at DIV 3. The media was replaced with half-fresh half-conditioned media after 3 h and the cells were used 4 days later (DIV 7). To validate knockdown, neurons were lysed in ice-cold lysis buffer (20 mM HEPES pH 7.4, 100 mM NaCl, 1% Triton X-100, 5 mM EDTA, HALT Protease Inhibitor), resolved by SDS–PAGE and processed for Western blotting using anti-ABCA1 rabbit polyclonal and anti-β-actin mouse monoclonal as a loading control, anti-ABCA7 rabbit polyclonal and anti-β-actin mouse monoclonal as a loading control.

### Immunostaining protocol

Cells were fixed in 4% paraformaldehyde (PFA) for 10 min at room temperature, washed twice in PBS with 0.1% Triton X-100 and blocked with PBS containing 2% bovine serum albumin and 0.2% Triton X-100 for 1 h at room temperature. Cells were incubated with primary antibody diluted in blocking buffer for 1 h at room temperature, washed three times in PBS with 0.1% Triton X-100 and incubated with secondary antibody diluted in blocking buffer for 1 h at room temperature. After washing, the cells were incubated with DAPI diluted in PBS for 10 min and mounted using DAKO fluorescence mounting media. For quantification, 10 images per coverslip were averaged.

### Confocal microscopy

Imaging was performed using the Laser Scanning Confocal Microscope (LSM) 900 with Airyscan 2 equipped with a plan-apochromat 40x oil objective (Zeiss, NA = 1.3) and 63x oil objective (Zeiss, NA = 1.4) and ZEN software (Zeiss). All imaging was performed using a 63x oil objective and SR-2Y mode unless otherwise specified.

### Lipid droplet assays

Neurons plated on PDL-coated glass coverslips were washed three times in warm PBS and treated with or without 50 μg/ml ApoE particles in HBSS for 4 h at 37°C. Cells were fixed in 4% PFA and stained with 5 μg/ml BODIPY 493/503 for 1 h at room temperature. The imaging field of view was selected using the DAPI channel to blind the experimenter to the BODIPY 493/503 channel. 10 images were averaged per coverslip with 1-2 cells per image. Maximum intensity projections of three-dimensional image stacks were generated. The images were thresholded using the Yen algorithm and the number of BD493-positive lipid droplets per nuclei were analyzed using Fiji-ImageJ software.

### Exogenous lipid release assays

Neurons were grown on PDL-coated 6-well plastic dishes. For fatty acid and cholesterol assays, neurons were loaded with 2 μM Red-C12 or 2 μM NBD cholesterol for 16 h in complete media. Cells were washed three times in warm PBS, incubated with fresh media for 1 h and treated with or without 50 μg/ml ApoE particles in HBSS for 4 h at 37°C. For lipid peroxidation assays, neurons were incubated with 2 μM BODIPY 581/591 C11 (BD-C11) for 1 h in complete media. Cells were washed three times in warm PBS and treated with or without DMSO, 10 μM Erastin, 10 μM Ferrostatin-1 (FS-1) or 50 μg/ml ApoE particles in HBSS for 4 h at 37°C. Neuron-conditioned media was collected and centrifuged at 16,000 x g for 15 min to remove dead cells or cell debris. The supernatant was analyzed for Red-C12 (excitation/emission 561/600 nm), NBD cholesterol (excitation/emission 465/535 nm) or BD-C11 (excitation/emission 488/530 and 568/619 nm) fluorescence using Synergy Mx Multi-Mode Microplate Reader (BioTek Instruments Inc.). Each treatment was analyzed in triplicate and averaged for quantification.

### Endogenous cholesterol release assays

Neurons were plated on 10-cm PDL-coated plastic plates. Neurons were washed twice in warm PBS and incubated in HBSS with or without 50 μg/ml ApoE particles for 6 h. Neuron-conditioned media was collected and centrifuged at 10,000 x g for 20 min at 4°C to remove dead cells or cell debris. The supernatant was transferred to ultra-4 centrifugal filter units with a 10 kDa cutoff and centrifuged at 4,000 x g for 25 min at 4°C to concentrate the media and total cholesterol content was analyzed using a cholesterol/cholesterol ester-glo assay kit according to the manufacturer’s protocol and Synergy Mx Multi-Mode Microplate Reader (BioTek Instruments Inc.).

### Endogenous unsaturated fatty acid release assays

Neurons were plated on 10-cm PDL-coated plastic plates. Neurons were washed twice in warm PBS and incubated in HBSS with or without DMSO, 10 μM Erastin, 10 μM Ferrostatin-1 (FS-1) or 50 μg/ml ApoE particles for 6 h. Neuron-conditioned media was collected and centrifuged at 10,000 x g for 20 min at 4°C to remove dead cells or cell debris. The supernatant was transferred to ultra-4 centrifugal filter units with a 10 kDa cutoff and centrifuged at 2,000 x g for 30 min at 4°C to concentrate the media and the lipid content (unsaturated fatty acids) was analyzed using a lipid assay kit (unsaturated fatty acids) according to the manufacturer’s protocol and Synergy Mx Multi-Mode Microplate Reader (BioTek Instruments Inc.). Each treatment was analyzed in triplicate and averaged for each biological replicate.

### Lipid peroxidation assays

For BD-C11 lipid peroxidation assay, neurons plated on PDL-coated glass coverslips were washed three times in warm PBS and incubated in complete media with or without DMSO, 100 μM Erastin or 50 μg/ml ApoE particles for 4 h with 2 μM BD-C11 added for the last 60 min. Cells were fixed and lipid peroxidation was determined by imaging oxidized sensor (excitation/emission - 490/565 nm) and non-oxidized sensor (excitation/emission - 570/700 nm). The imaging field of view was selected using the DAPI channel to blind the experimenter to the BD-C11 channels. 10 images were averaged per coverslip with 1 cell per image. The relative lipid peroxidation was indicated as the ratio of green fluorescence intensity over red fluorescence intensity using Fiji-ImageJ. For lipid hydroperoxide assay, neurons were grown on PDL-coated 6-well plastic dishes. Neurons were washed three times in warm PBS and incubated in HBSS or complete media with or without NaOH, DMSO, 2 μM Aβ42, 10 μM Pitstop 2 or 50 μg/ml ApoE particles for 6 h. Neuron-conditioned HBSS was collected and neurons used for the assay were lysed in ice-cold lysis buffer (20 mM HEPES pH 7.4, 100 mM NaCl, 1% Triton X-100, 5 mM EDTA, HALT Protease Inhibitor). For tissue, ∼50-150 mg hippocampal tissue extracted as described below under ‘ex vivo intact hippocampi incubation’ was homogenized in 550 μL HPLC-grade water using a pellet pestle. Lipid hydroperoxides were extracted from neuronal lysates, neuron-conditioned HBSS, and tissue homogenates and analyzed using a lipid hydroperoxide assay kit according to the manufacturer’s protocol and Synergy Mx Multi-Mode Microplate Reader (BioTek Instruments Inc.). Each treatment was analyzed in triplicate and averaged for quantification. Since the background signal is increased when lipid hydroperoxides are extracted in polypropylene tubes as performed in this study, lipid hydroperoxides were reported as relative values. Cell culture assays were normalized to the control treatment for each independent assay, while ex vivo tissue assays were normalized to the tissue weight for each hippocampus.

### Total cell death assays

Neurons were plated on 96-well PDL-coated plastic plates. Neurons were washed twice in warm PBS and treated with or without NaOH, DMSO, 2 μM Aβ42, 100 μM Erastin or 50 μg/ml ApoE particles in complete media for 12 h. Neurons were washed twice in warm PBS and incubated with 4 μM Ethidium Homodimer-1 in PBS for 30 min at 37°C to label dead cells, washed twice in PBS and fluorescence was measured using Synergy Mx Multi-Mode Microplate Reader (BioTek Instruments Inc.). Three wells per treatment were analyzed and averaged for quantification.

### Lysosome accumulation assays

Neurons plated on PDL-coated glass coverslips were washed twice in warm PBS and treated with or without 50 μg/ml ApoE particles in HBSS for 4 h at 37°C. Neurons were fixed in 4% PFA and immunostained for LAMP1 to label lysosomes as described above. Imaging was performed using a 40x oil objective and SR-2Y mode. The imaging field of view was selected using the DAPI channel to blind the experimenter to the LAMP1 channel. 10 images were averaged per coverslip with 1-2 cells per image. First, maximum intensity projections of three-dimensional image stacks of LAMP1 staining were obtained. To remove background, a Gaussian blur was applied to a duplicate image and subtracted from the original image. The images were thresholded using the Yen algorithm and the number of LAMP1-positive puncta were analyzed using Fiji-ImageJ software.

### Lysosome function assays

For cathepsin D assay, neurons were grown on 6-well plastic dishes. Neurons were washed twice in warm PBS and incubated in HBSS or complete media with or without DMSO, 10 μM Erastin (HBSS), 100 μM Erastin (CM) or 50 μg/ml ApoE particles for 6 h. For tissue, the hippocampi were homogenized in HPLC-grade water. Neuronal lysates and tissue homogenates were analyzed using a cathepsin D activity assay kit according to the manufacturer’s protocol and Synergy Mx Multi-Mode Microplate Reader (BioTek Instruments Inc.). Each treatment was analyzed in triplicate and averaged for quantification. Cell culture assays were normalized to the control treatment for each independent assay, while ex vivo tissue assays were normalized to the protein levels for each hippocampus as calculated by a BCA assay. For DQ-BSA assay, neurons plated on PDL-coated glass coverslips were washed twice in warm PBS and incubated in HBSS or complete media with or without DMSO, 10 μM Erastin (HBSS), 100 μM Erastin (CM) or 50 μg/ml ApoE particles for 4 h with 10 μg/ml DQ-BSA added for the last 60 min. Cells were fixed in 4% PFA. Imaging and quantification were performed blind. The imaging field of view was selected using the DAPI channel. 10 images were averaged per coverslip with 1 cell per image. Mean values were obtained for DQ Red BSA fluorescence intensity using Fiji-ImageJ software.

### Iron accumulation and release assays

Neurons were grown on PDL-coated 6-well plastic dishes. Neurons were washed twice in warm PBS and incubated in HBSS with or without 50 μg/ml ApoE particles for 6 h. Total iron (Fe^2+^ and Fe^3+^) from neuronal lysates and neuron-conditioned media was measured using an iron assay kit according to the manufacturer’s protocol and Synergy Mx Multi-Mode Microplate Reader (BioTek Instruments Inc.). Each treatment was analyzed in triplicate and averaged for quantification.

### ApoE particle internalization assay

Neurons were grown on PDL-coated 6-well plastic dishes. Neurons were washed twice in warm PBS and treated with or without 25 μg/ml ApoE particles in HBSS for 0, 15 or 30 min. Neurons were washed twice in ice-cold acid wash (0.5 M NaCl and 0.2 M acetic acid pH 2.5), followed by two washes with ice-cold PBS. Neurons were lysed in ice-cold lysis buffer, resolved by SDS-PAGE and processed for Western Blotting using anti-ApoE goat polyclonal and anti-β-actin mouse monoclonal as a loading control.

### Ex vivo intact hippocampi incubation

Human ApoE2, ApoE3 or ApoE4 targeted replacement rats (P9-P10) were decapitated and the brain was rapidly removed from the skull and placed in ice-cold high sucrose artificial cerebrospinal fluid (aCSF) solution (252 mM sucrose, 3 mM KCl, 2 mM MgSO_4_, 24 mM NaHCO_3_, 1.25 mM NaH_2_PO_4_, 1.2 mM CaCl_2_ and 10 mM glucose) bubbled with carbogen (95% O_2_ and 5% CO_2_). The hippocampi were removed from the brain and transferred to fresh ice-cold high sucrose aCSF with or without 50 μg/mL ApoE particles. Following dissection, the hippocampus was left at room temperature in the high sucrose aCSF bubbled with carbogen for 30 min to rest. The hippocampi were then transferred to a new chamber and continuously perfused with aCSF with or without 50 μg/mL ApoE particles for 5 h via a gravity fed perfusion system and maintained at 30-32°C. For the first 30 min, the preparation was perfused with control aCSF (125 mM NaCl, 5 mM KCl, 2 mM MgSO_4_, 24 mM NaHCO_3_, 1.25 mM NaH_2_PO_4_, 2 mM CaCl_2_ and 10 mM glucose pH 7.4), followed by 4.5 h with low magnesium high potassium aCSF (125 mM NaCl, 8 mM KCl, 0.5 mM MgSO_4_, 24 mM NaHCO_3_, 1.25 mM NaH_2_PO_4_, 2 mM CaCl_2_ and 10 mM glucose pH 7.4). Following treatment, the hippocampus was flash frozen in liquid nitrogen.

### Local field potential electrophysiological recordings

ApoE3 or ApoE4 hippocampi were removed from the brain and allowed to rest in oxygenated high sucrose aCSF with or without ApoE particles as described above. For recording, the hippocampus was transferred to the recording chamber containing control aCSF solution (125 mM NaCl, 5 mM KCl, 2 mM MgSO_4_, 24 mM NaHCO_3_, 1.25 mM NaH_2_PO_4_, 2 mM CaCl_2_ and 10 mM glucose pH 7.4) with or without 50 μg/ml ApoE2 or ApoE3Ch particles and the preparation was continuously perfused with control aCSF (15 mL/min) for 30 min via a gravity fed perfusion system and maintained at 30-32°C. Local field potentials were recorded using a tungsten electrode placed in the subiculum region. Signals were recorded using an OpenEphys acquisition board and software and sampled at 10KHz. The signals were referenced to the bath medium and connected to the ground. The hippocampi were then continuously perfused with low magnesium high potassium (LMHK) aCSF (15 mL/min; 125 mM NaCl, 8 mM KCl, 0.5 mM MgSO_4_, 24 mM NaHCO_3_, 1.25 mM NaH_2_PO_4_, 2 mM CaCl_2_ and 10 mM glucose pH 7.4) with or without 50 μg/ml ApoE2 or ApoE3Ch particles for 4.5 h. Following the recording, the hippocampus was flash frozen in liquid nitrogen.

### Electrophysiology analysis

Local field potential (LFP) data were processed in Matlab 2018b. First, high amplitude noise events were removed from the data by manual inspection. Data were low pass filtered below 30Hz. The number of LFP peaks were identified by counting high amplitude positive going LFP peaks that exceeded five standard deviations above the mean value of the baseline data period (prior to low magnesium high potassium aCSF bath application). To detect LFP peaks we used the ‘findpeaks’ function in Matlab and used a minimum peak distance of 500s. This ensured multiple peaks within the same LFP burst were not counted. The overall amplitude of the LFP signal was calculated by filtering the full LFP timeseries (lowpass 30Hz), and then taking the absolute value of the filtered data and performing a 5s moving average smoothing of the data. The data were normalized to baseline values for both the number of LFP events and the amplitude of the LFP changes.

### Sex determination in rats

Sex determination of rat tissues was performed as previously described (Dhakal and Soares^44^). Briefly, genomic DNA was isolated from P9-P10 rat cortical tissue using the Extract-N-Amp tissue PCR kit according to the manufacturer’s protocol. A PCR reaction mix containing the primers was prepared and amplification was performed using the BioRad T100 Thermal Cycler. PCR products were resolved by agrose gel electrophoresis and UV-light imaging was performed using a BioRad Gel Doc EZ Imager. The PCR reaction generated 692 base pair (bp) X chromosome-specific amplicons and 250 bp Y chromosome-specific amplicons.

## STATISTICAL ANALYSES

Datasets were assembled in Microsoft Excel 365 (Microsoft Corp.). Statistical analysis and graphing were performed using Graphpad Prism 10. All graphs are depicted as Superplots where independent replicates are shown in large shapes, and the corresponding technical replicants shown as small shapes of the same color. Statistical test used and p-values can be found in the figures and/or figure legends.

---

**Figure S1.**
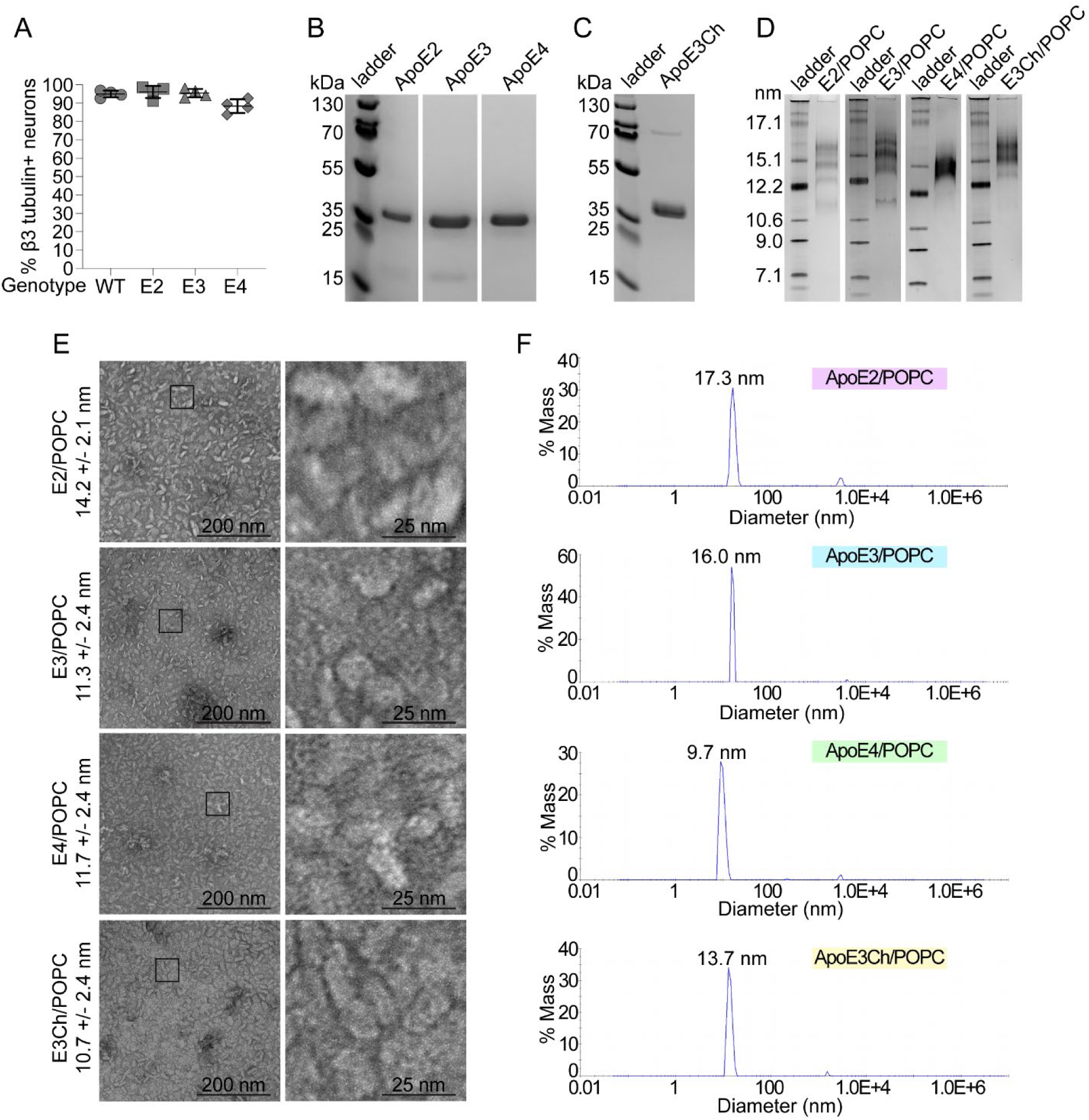
Validation of ApoE particles used to study lipid transport, related to Figure 1 (A) Neuronal cultures grown were fixed and immunostained for neuron-specific β3-tubulin and the percentage of β3-tubulin-positive neurons was quantified. n = 4 independent experiments; mean ± SD. (B and C) Purified ApoE proteins were analyzed by reducing sodium dodecyl sulfate - polyacrylamide gel electrophoresis (PAGE) and Coomassie staining. (D) ApoE particles were analyzed by native PAGE and Coomassie staining for size. (E) ApoE particles imaged by transmission electron microscopy. Boxed areas magnified in bottom panels. Scale bars are 200 nm and 25 nm. (F) ApoE particle diameters were assessed by dynamic light scattering. Related to Table S1. For all graphs independent replicates are in large shapes and technical replicates in small shapes; *p < 0.05, **p < 0.01, ***p < 0.001, ****p < 0.0001.

**Figure S2.**
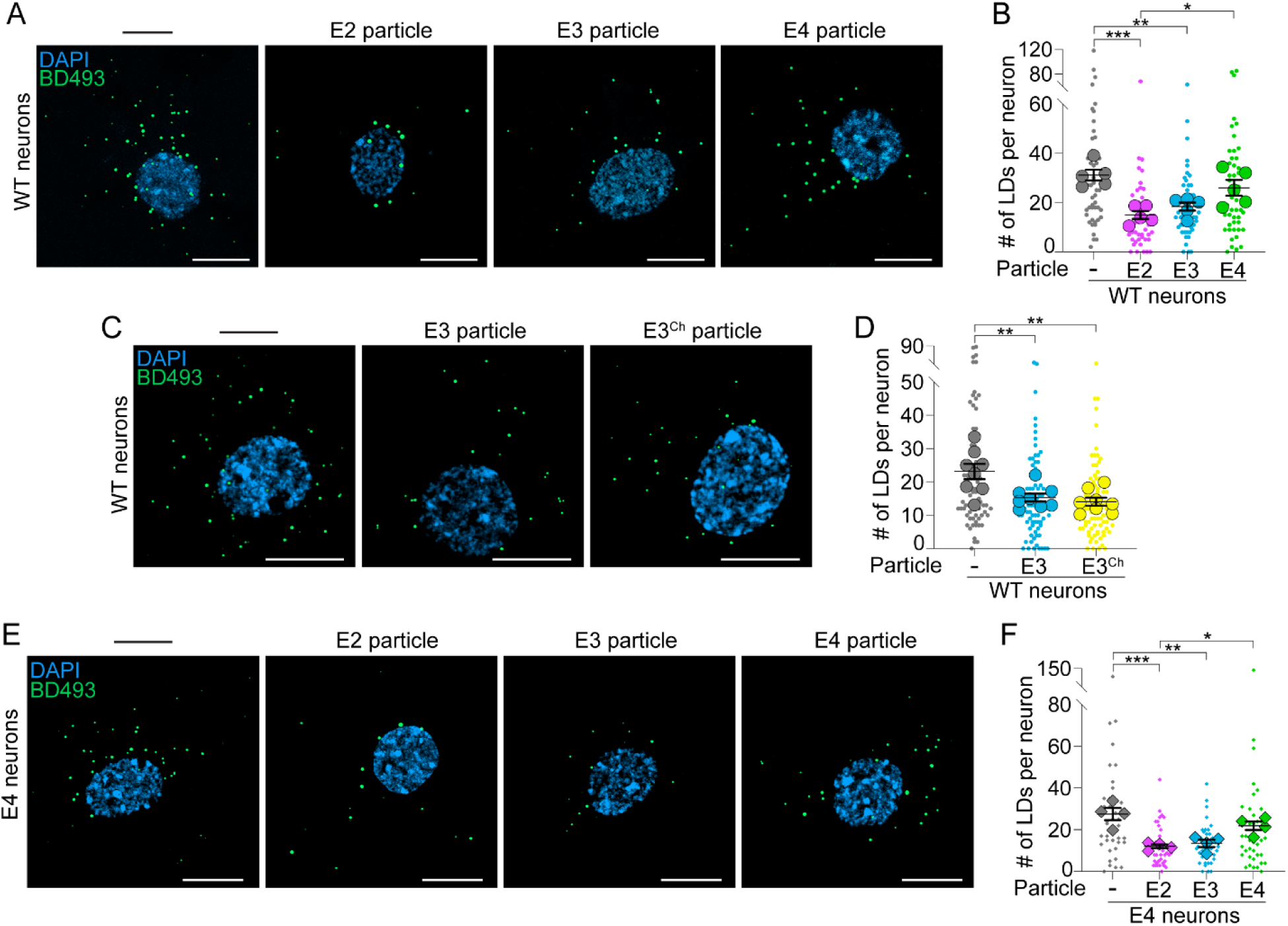
ApoE variants differentially regulate fatty acid accumulation in neurons, related to Figure 1 (A and B) Airyscan images of wild-type (WT) neurons following ApoE2, ApoE3 or ApoE4 particle treatment in HBSS stained for BD493 positive lipid droplets (LDs). n = 5 independent experiments; mean ± SEM. One-way ANOVA with Tukey’s post-test. Scale bars are 10 μm. (C and D) Airyscan images of WT neurons following ApoE3 or ApoE3^Ch^ particle treatment in HBSS stained for BD493 positive LDs. n = 8 independent experiments; mean ± SEM. One-way ANOVA with Tukey’s post-test. Scale bars are 10 μm. (E and F) Airyscan images of ApoE4 neurons following ApoE2, ApoE3 or ApoE4 particle treatment in HBSS stained for BD493 positive LDs. n = 4 independent experiments; mean ± SEM. One-way ANOVA with Tukey’s post-test. Scale bars are 10 μm. All images displayed as maximum intensity projections. For graphs, independent replicates are large shapes and technical replicates in small shapes; *p < 0.05, **p < 0.01, ***p < 0.001.

**Figure S3.**
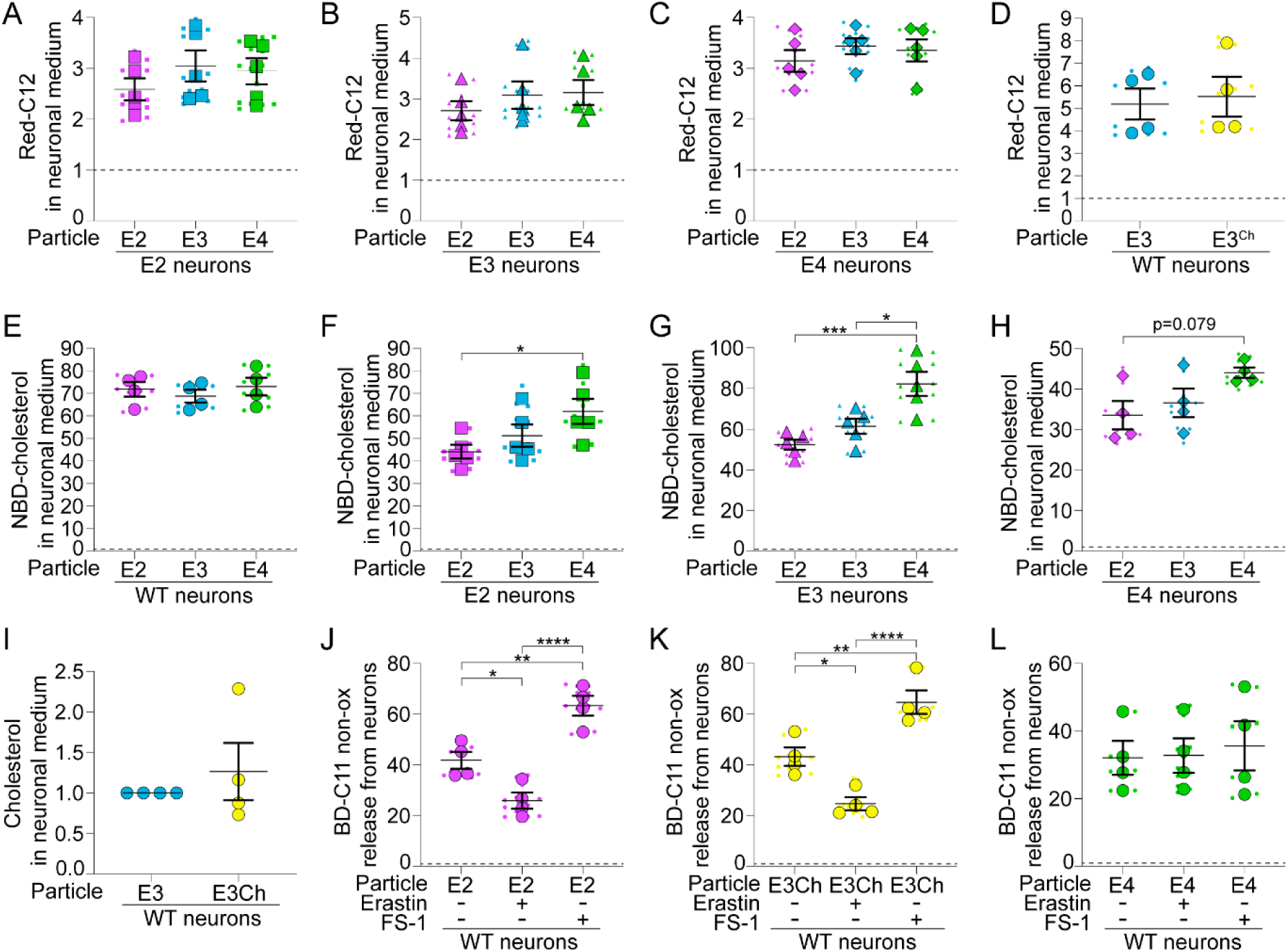
ApoE isoforms differentially regulate lipid release from neurons, related to Figure 1 (A–C) ApoE targeted replacement neuron-conditioned HBSS following ApoE2, ApoE3 or ApoE4 particle treatment analyzed for Red-C12 fluorescence and normalized to non-particle treated control neurons (dashed line). n = 5 independent experiments; mean ± SEM. One-way ANOVA with Tukey’s post-test. (D) WT neuron-conditioned HBSS following ApoE3 or ApoE3^Ch^ particle treatment analyzed for Red-C12 fluorescence and normalized to non-particle treated control neurons (dashed line). n = 4 independent experiments; mean ± SEM. Two-tailed Student’s t test. (E-H) WT or ApoE targeted replacement neuron-conditioned HBSS following ApoE2, ApoE3 or ApoE4 particle treatment analyzed for NBD-cholesterol fluorescence and normalized to non-particle treated control neurons (dashed line). n = 4-5 independent experiments; mean ± SEM. One-way ANOVA with Tukey’s post-test. (I) WT neuron-conditioned HBSS following ApoE3 or ApoE3^Ch^ particle treatment analyzed for endogenous cholesterol and normalized to ApoE3 particle treated neurons. n = 4 independent experiments; mean ± SEM. One sample t test with Bonferroni correction. (J-L) WT neuron-conditioned HBSS following ApoE2, ApoE3^Ch^ or ApoE4 particle treatment with or without erastin or ferrostatin-1 (FS-1) in HBSS, analyzed for non-peroxidated lipids (BD-C11 non-ox) and normalized to DMSO treated control neurons (dashed line). n = 4 independent experiments; mean ± SEM. One-way ANOVA with Tukey’s post-test. For all graphs independent replicates are in large shapes and technical replicates in small shapes; *p < 0.05, **p < 0.01, ***p < 0.001, ****p < 0.0001.

**Figure S4.**
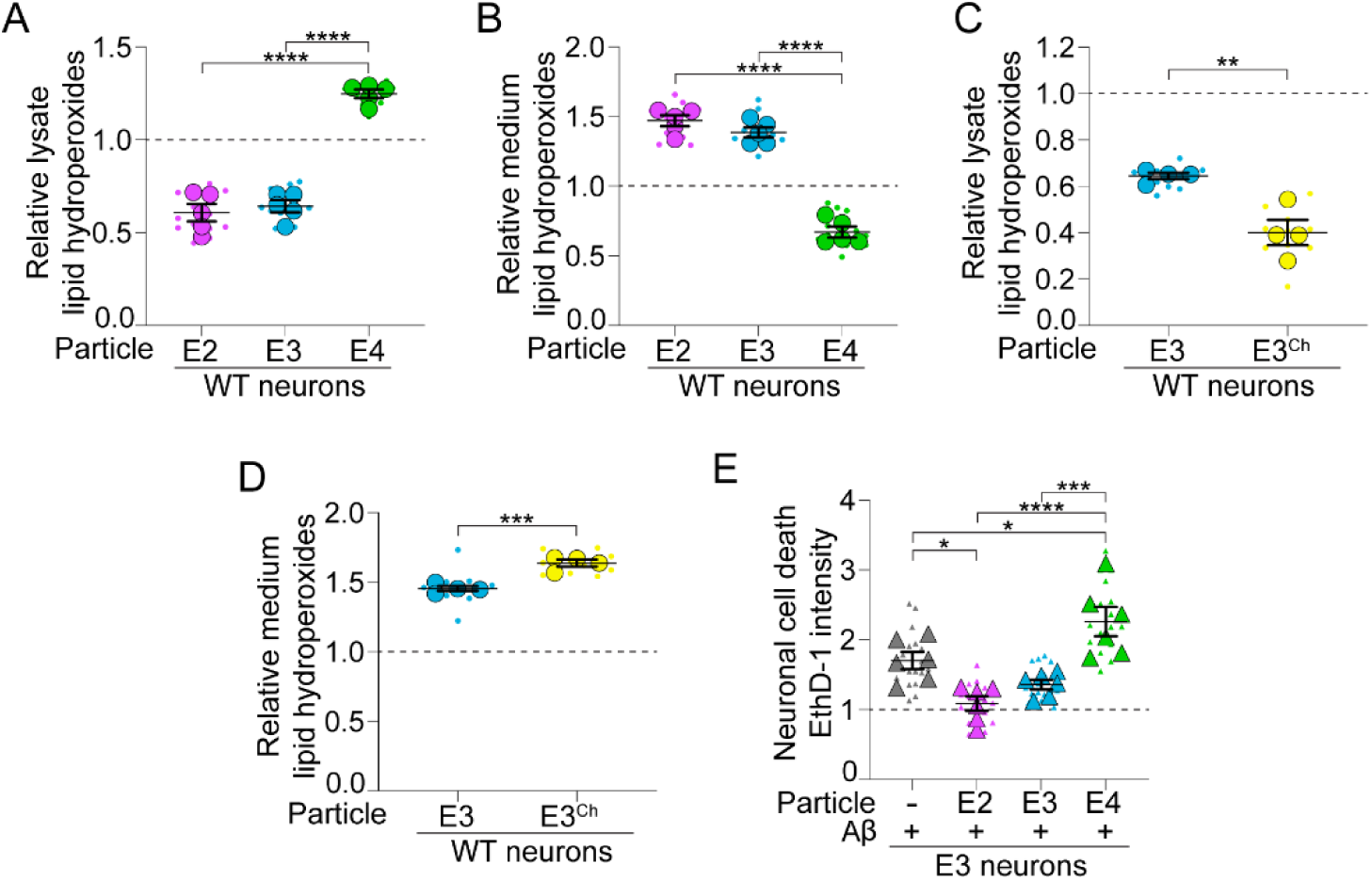
ApoE2 and ApoE3Ch particles protect neurons from cell death, related to Figure 2 (A and B) WT neuronal lysates (A) and conditioned HBSS (B) following ApoE2, ApoE3 or ApoE4 particle treatment were analyzed for lipid hydroperoxides and normalized to non-particle treated control neurons (dashed line). n = 5 independent experiments; mean ± SEM. One-way ANOVA with Tukey’s post-test. (C and D) WT neuronal lysates (C) and neuron-conditioned HBSS (D) following ApoE3 or ApoE3^Ch^ particle treatment were analyzed for lipid hydroperoxides and normalized to non-particle treated control neurons (dashed line). n = 4 independent experiments; mean ± SEM. Two-tailed Student’s t test. (E) ApoE3 neurons following Aβ, ApoE2, ApoE3 or ApoE4 particle treatment in media were analyzed for ethidium homodimer-1 (EthD-1) intensity and normalized to NaOH treated control neurons (dashed line). n = 6 independent experiments; mean ± SEM. One-way ANOVA with Tukey’s post-test. For all graphs independent replicates are in large shapes and technical replicates in small shapes; *p < 0.05, **p < 0.01, ***p < 0.001, ****p < 0.0001.

**Figure S5.**
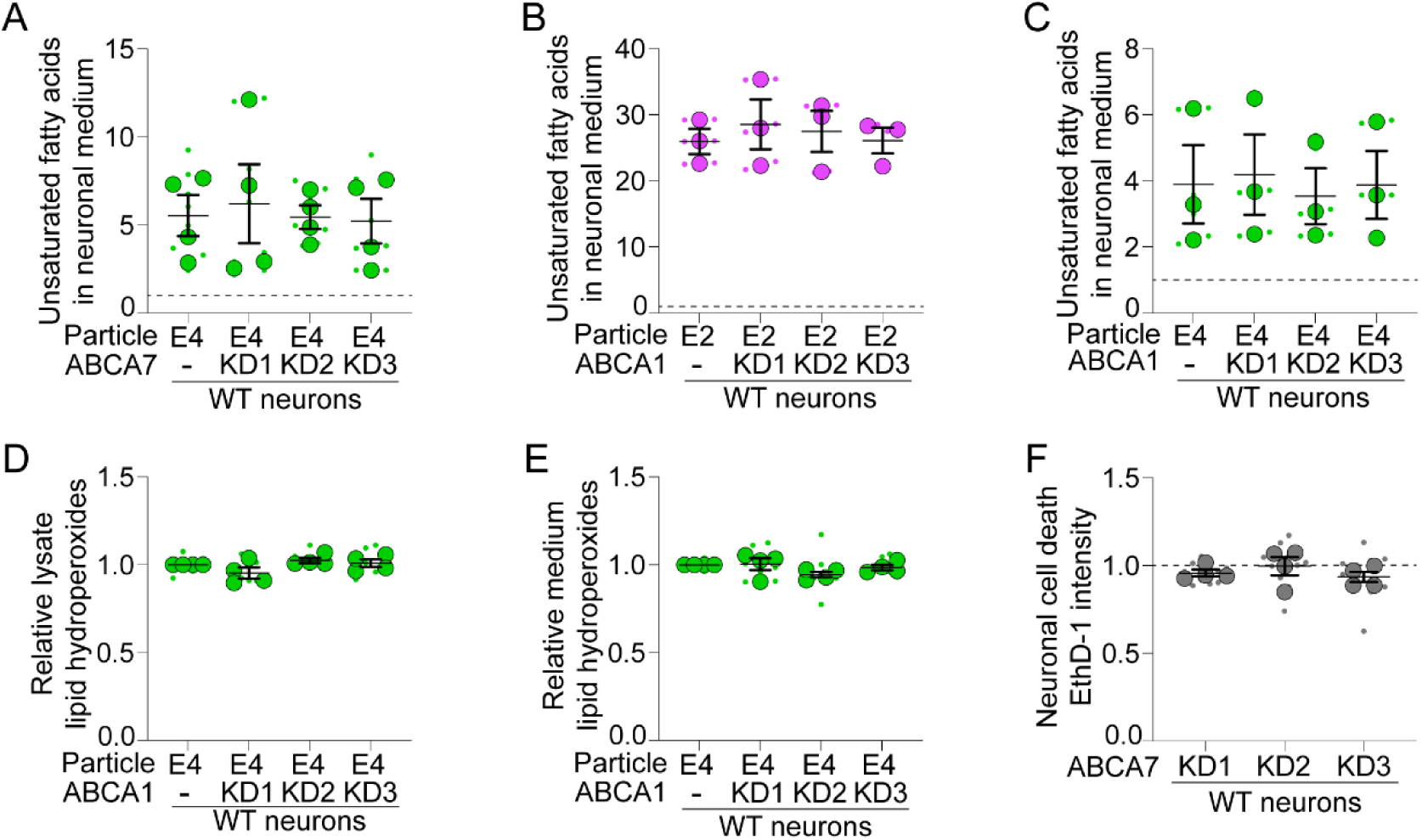
ABCA1 does not affect ApoE-mediated peroxidated lipid efflux from neurons, related to Figure 3 (A) WT neurons were treated with ABCA7 or non-targeting shRNAmiR. After ApoE4 particle treatment conditioned HBSS was analyzed for endogenous unsaturated fatty acids. ABCA7 knockdown neurons were normalized to non-targeting shRNAmiR non-particle treated control neurons (dashed line). n = 4 independent experiments; mean ± SEM. One-way ANOVA with Tukey’s post-test. (B and C) WT neurons were treated with ABCA1 or non-targeting shRNAmiR. After ApoE2 or ApoE4 particle treatment conditioned HBSS was analyzed for endogenous unsaturated fatty acids. ABCA1 knockdown neurons were normalized to non-targeting shRNAmiR non-particle treated control neurons (dashed line). n = 3 independent experiments; mean ± SEM. One-way ANOVA with Tukey’s post-test. (D and E) WT neurons were treated with ABCA1 or non-targeting shRNAmiR. After ApoE4 particle treatment lysates (D) and conditioned HBSS (E) were analyzed for lipid hydroperoxides. ABCA1 knockdown neurons were normalized to non-targeting shRNAmiR ApoE4 particle treated control neurons. n = 4 independent experiments; mean ± SEM. One sample t test with Bonferroni correction. (F) WT neurons treated with ABCA7 or non-targeting shRNAmiR analyzed for ethidium homodimer-1 (EthD-1) intensity and normalized to non-targeting control neurons (dashed line). n = 4 independent experiments; mean ± SEM. One-way ANOVA with Tukey’s post-test. For all graphs independent replicates are in large shapes and technical replicates in small shapes; *p < 0.05, **p < 0.01, ***p < 0.001, ****p < 0.0001.

**Figure S6.**
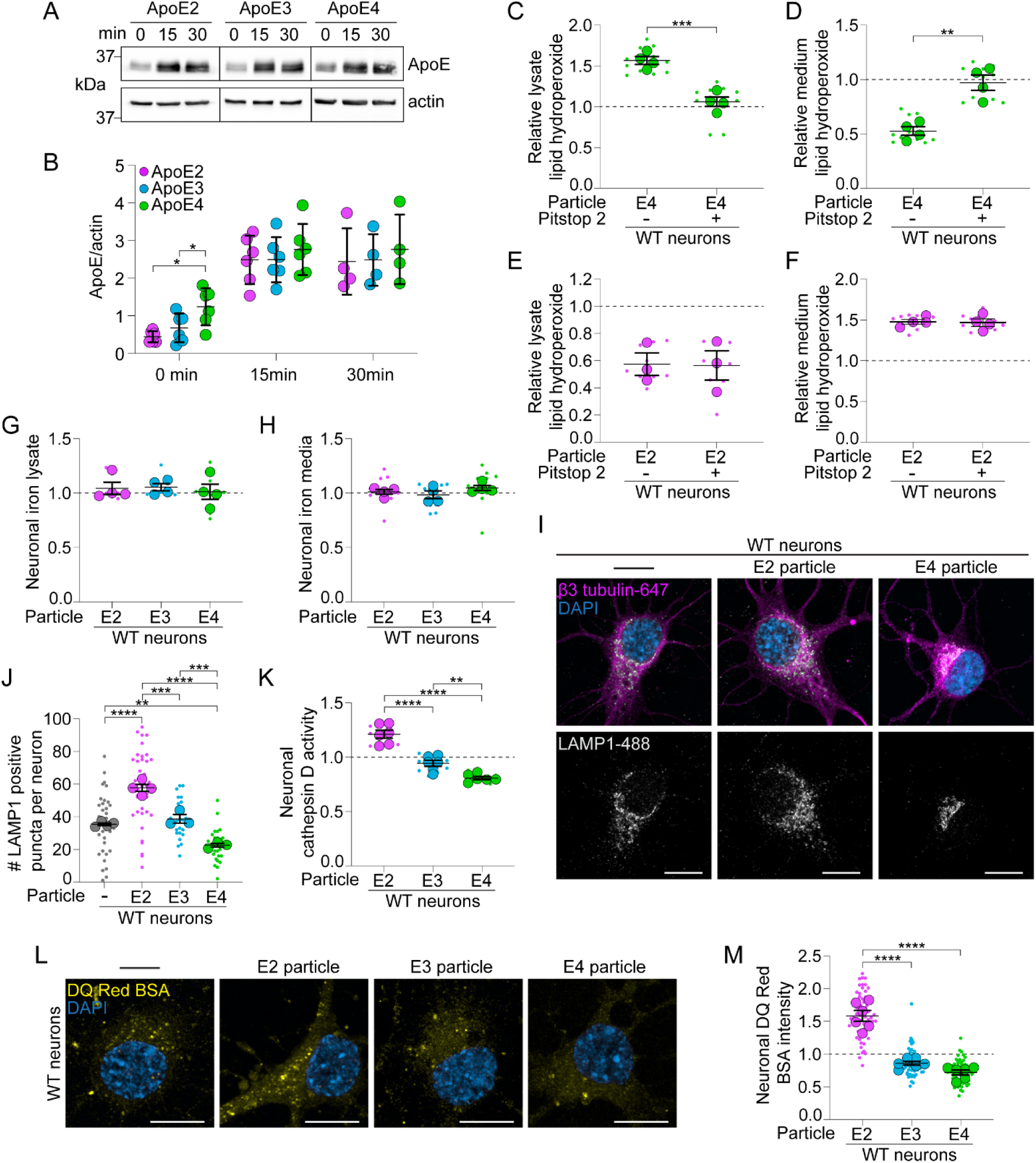
ApoE4 induced toxicity is rescued by ApoE2 particles, related to Figure 4 (A and B) WT neurons following ApoE2, ApoE3 or ApoE4 particle treatment in HBSS were acid washed and analyzed by Western blot for ApoE and β-actin. n = 4-6 independent experiments; mean ± SD. One-way ANOVA with Tukey’s post-test. (C and D) WT neuronal lysates (C) and neuron-conditioned HBSS (D) following ApoE4 particle treatment with or without Pitstop 2 were analyzed for lipid hydroperoxides and normalized to DMSO treated control neurons (dashed line). n = 4 independent experiments; mean ± SEM. Two-tailed Student’s t test. (E and F) WT neuronal lysates (E) and neuron-conditioned HBSS (F) following ApoE2 particle treatment with or without Pitstop 2 were analyzed for lipid hydroperoxides and normalized to DMSO treated control neurons (dashed line). n = 3-4 independent experiments; mean ± SEM. Two-tailed Student’s t test. (G and H) WT neuronal lysates (G) and neuron-conditioned HBSS (H) following ApoE2, ApoE3 or ApoE4 particle treatment were analyzed for iron and normalized to non-particle treated control neurons (dashed line). n = 4 independent experiments; mean ± SEM. One-way ANOVA with Tukey’s post-test. (I and J) Airyscan maximum intensity projection images of WT neurons following ApoE2, ApoE3 or ApoE4 particle treatment in HBSS, immunostained for LAMP1 and the number of LAMP1 positive puncta were quantified. n = 3-4 independent experiments; mean ± SEM. One-way ANOVA with Tukey’s post-test. Scale bars are 10 μm. (K) WT neurons following ApoE2, ApoE3 or ApoE4 particle treatment in HBSS analyzed for cathepsin D activity and normalized to non-particle treated control neurons (dashed line). n = 6 independent experiments. mean ± SEM. One-way ANOVA with Tukey’s post-test. (L and M) Airyscan maximum intensity projection images of WT neurons following ApoE2, ApoE3 or ApoE4 particle treatment in HBSS, labeled with DQ Red BSA and normalized to non-particle treated control neurons (dashed line). n = 6 independent experiments; mean ± SEM. One-way ANOVA with Tukey’s post-test. Scale bars are 10 μm. For all graphs independent replicates are in large shapes and technical replicates in small shapes; *p < 0.05, **p < 0.01, ***p < 0.001, ****p < 0.0001.

